# Drought stress modulates the molecular response of Arabidopsis plants to root-knot nematode infection

**DOI:** 10.1101/2025.06.05.658137

**Authors:** Ahmed Refaiy, Catherine J Lilley, Nicky J Atkinson, Peter E Urwin

**Author notes:** **Corresponding authors:** Ahmed Refaiy; Peter E Urwin.

## Abstract

Plants are exposed to multiple concurrent stresses under field conditions that can include a combination of biotic and abiotic factors. Drought stress and plant parasitic nematodes are among the most damaging stress factors limiting plant growth and production. Plants have developed complex and specific responses to combined biotic and abiotic stresses, which differ from their responses to individual stresses. Therefore, there is an imperative need to understand plant responses to multiple stresses as a central avenue for development of robust plants with the ability to sustain growth and crop production in times of global climate change. RNA-Seq analysis was performed on Arabidopsis plants to investigate the transcriptomic responses to infection with the root-knot nematode *Meloidogyne incognita* and drought stress either individually or concurrently. Arabidopsis plants activated a unique suite of genes in response to the joint stress, which significantly differed from the response to either individual stress. Among these differentially expressed genes, *Azelaic Acid Induced 1 (AZI1), small auxin upregulated RNA 71 (SAUR71),* and *Disease Related Nonspecific Lipid Transfer Protein 1 (DRN1)* may play important roles in plant responses to concurrent drought stress and root-knot nematode infection. The expression of *AZI1*, which is involved in priming defences via systemic acquired resistance, was uniquely upregulated in Arabidopsis leaves in response to the combined stress. *SAUR71,* which is a member of the largest family of primary auxin response genes, was induced solely in response to combined stress. In contrast, the expression of *DRN1,* a member of the non-specific lipid-transfer protein family, was strongly suppressed in Arabidopsis roots by combined drought stress and nematode infection.

## Introduction

Plants in natural environments are subjected to multiple concurrent stresses including a combination of several abiotic and biotic stress factors. Unfortunately, many studies have focused on the plant response to individual stresses that are unlikely to occur alone in field conditions (Mittler, 2006; Pandey *et al*., 2017). Plant responses to multiple stresses are significantly different from their responses to individual stress factors, but are not merely additive (Atkinson & Urwin, 2012; Rivero *et al*., 2014; Suzuki *et al*., 2014). Rather than produce a stock response to each pathogen or abiotic stress, plants subjected to a combination of stresses induce a novel programme of gene expression, activating transcripts that are not induced by either stress individually (Rizhsky *et al.,* 2002; Atkinson *et al.,* 2013; Rasmussen *et al.,* 2013). Concurrent biotic and abiotic stresses not only modify the plant responses, but also modulate the growth and development of pathogens and affect disease severity (Triky-Dotan *et al*., 2005; Al-Sadi *et al*., 2010).

Stress causes numerous disorders in plants, which adversely affect their growth and yield production. Therefore, plants induce different short- and long-term adaptive responses, tailored to the particular stress, to cope with these effects. For example, stomatal closure is considered an important strategy in plants under drought stress conditions to decrease stomatal conductance and prevent transpiration water loss (Martin-StPaul *et al.,* 2017). On the other hand, plants grown under heat stress conditions tend to open their stomata to increase stomatal conductance, which consequently increases transpiration rates as a strategy for leaf cooling (Ameye *et al.,* 2012). Under combined drought and heat stresses, the effect of drought stress was dominant in leaves which displayed stomatal closure and decreased stomatal conductance. This elevated the leaf temperatures of plants grown under combined stresses by 2-3°C compared to plants grown under heat stress only (Rizhsky *et al.,* 2002). In contrast, the heat stress response was dominant in flowers, which opened their stomata, decreasing flower temperature by 2-3°C (Sinha *et al.,* 2022). This highlights the tissue-specific differences of plant responses to multiple environmental stresses.

Many environmental factors influence plant-pathogen interactions, which increases the complexity of plant responses and tolerance to pathogens (Atkinson & Urwin, 2012; Omae & Tsuda, 2022). High temperature generally increases the spread of plant pathogens and facilitates their infections (Haverkort & Verhagen, 2008). It reduces the biosynthesis of salicylic acid in plants and suppresses their basal immunity against pathogens (Kim *et al.,* 2022). For example, Zacheo and Bleve-Zacho (1984) reported that elevated temperature significantly increased the total number of nematodes and reduced the resistance of tomato plants infected with the root-knot nematode *Meloidogyne incognita*. In addition, high temperatures above 28°C cause failure of the resistance imparted by the *Mi-1* gene in tomato plants, which increases the plant susceptibility to *M. incognita* infection (Dropkin, 1969; de Carvalho *et al.,* 2015). Drought stress also affects plant pathogen resistance. It increases the susceptibility of rice plants to the blast fungus *Magnaporthe oryzae* by inhibiting basal immunity and thus increasing disease symptoms (Bidzinski *et al.,* 2016). Multifactorial stress leads to severe growth reduction compared to individual stresses. Plants subjected to combined drought, heat, and Turnip mosaic virus showed pronounced reductions in growth compared to either individual stress alone. The combined drought and heat stress inhibited defence responses and increased the susceptibility to virus infections (Prasch & Sonnewald, 2013).

The combined stress caused by concurrent drought and plant parasitic nematode infection represents a major threat to plant growth and production (Atkinson & Urwin, 2012). Drought stress can affect both the density and composition of plant parasitic nematode populations (Coyne *et al.,* 2001) and influences plant responses to nematode infection (Atkinson *et al.,* 2013). The detrimental effects of parasitic nematodes on plant growth are typically more severe during periods of reduced water availability. Water deficit increased the damage suffered by rice plants due to cyst nematodes with the combined stress significantly decreasing leaf area, and both leaf and root dry weight (Audebert *et al.,* 2000). Yield loss in rice plants associated with micronutrient deficiencies was higher during the dry season due to the interaction between abiotic stress and root-knot nematodes (Kreye *et al.,* 2009). Similarly, the combination of drought stress and cyst nematode infection affects potato plant growth and development with plants responding differently compared to each individual stress (Fasan & Haverkort, 1991). In addition to reducing yield quantity, concurrent drought stress and nematode infection can also affect the quality and composition of crop yield. For example, tomato fruits from drought-stressed plants that were simultaneously infected with root-knot nematodes had significantly altered levels of carotenoids, sugars, and phenolic compounds (Atkinson *et al.,* 2011).

The molecular responses of plants to combined drought stress and nematode infection have been studied less frequently. Microarray analysis was previously used to investigate gene expression in leaves and roots of Arabidopsis plants in response to simultaneous drought stress and cyst nematode, *Heterodera schachtii*, infection (Atkinson *et al.,* 2013). That work uncovered a unique signature of transcriptional responses specific to the dual stress, despite a relatively low impact of the nematode infection alone. In the current work, we used a similar system to analyse the effect of acute drought stress on Arabidopsis plants infected with the root-knot nematode *M. incognita*. This sedentary endoparasite establishes a permanent feeding site in the vascular tissue of host plant roots, associated with morphological changes to root tissue, including the characteristic swelling or ‘knot’ that develops around nematode feeding cells (Escobar *et al*., 2015). The greater impacts of this nematode on root architecture, function and water relations may result in a more pronounced molecular response when infection is combined with drought stress.

## Material and methods

### RNA-seq analysis of Arabidopsis transcriptomic responses

RNA sequencing (RNA-seq) was used to identify the genes that differ significantly in expression in response to concurrent root-knot nematode infection and acute drought stress compared to either treatment alone. Seeds of *Arabidopsis thaliana* Col-0 were sterilized in 20% bleach for 20 minutes followed by five washes in sterile deionised water. The sterilized seeds were kept in water at 4 °C for 2 days for stratification, then they were grown vertically in square petri dishes on one-half strength MS basal medium with vitamins (Murashige & Skoog, 1962) containing 10 g/l plant agar (Duchefa) and 10 g/l sucrose. Three seeds were sown in each 10 cm square plate, approximately 2 cm from the top edge, then the plates were placed vertically in a Sanyo growth cabinet at 24 °C under 16 h-light and 8 h-dark regime.

After 18 days, half of the seedlings were infected with 2^nd^-stage juvenile (J2) nematodes of *M. incognita*. A total of 300 J2s were applied to each plant at 5 infection points distributed through the roots. A small piece of sterile GF/A microfibre filter paper was placed over each infection point to facilitate penetration by the J2s. These were removed under sterile conditions after 48 hours. At 28 days post sowing, half of the nematode-infected plants, and half of the uninfected plants were subjected to acute drought stress. The plants were removed from the agar and subjected to a clean flow of air in a flow hood for 15 minutes which induced a loss of about 10-15% of their fresh weight as described by Seki *et al*. (2002). The plants were returned to the agar for 30 minutes before harvesting the samples. Control plants were not exposed to either nematodes or drought stress. The control and nematode infection-only plants were transferred to fresh plates for 30 minutes to reproduce the physical handling experienced by the air-dried plants. Entire leaf and root tissues were harvested separately from each plant. Tissue from three plants was pooled to form a biological replicate, with three replicates per treatment for each tissue type. RNA was extracted from the leaves and roots (RNeasy Plant Kit, Qiagen). The quality of the extracted RNA was determined by the 2100 Expert Bioanalyser (Agilent) before RNA Sequencing was conducted by Genewiz following polyA selection, using the Illumina NovaSeq™ platform to generate 150 bp paired-end reads.

### RNA-seq data analysis

RNA-seq data analysis was performed using the Galaxy web service (https://usegalaxy.org/). Trimmomatic was performed to remove adapters from the paired-end reads. Reads were mapped onto the reference genome of *Arabidopsis thaliana* TAIR.10 using HISAT2. The read quantification was performed using HTSeq-count. DESeq2 was used to determine the differential expression and the statistical significance. The genes were considered to be differentially expressed if they were significantly different from the control where FDR < 0.05.

### Gene Ontology annotation

Gene ontology (GO) of the differentially expressed genes (DEGs) was analysed by the Arabidopsis Gene Ontology tool available at (https://www.arabidopsis.org/tools/bulk/go/index.jsp). The distributions of the DEGs among the GO terms biological process, molecular function, and subcellular location in the different treatments were compared to the whole genome distribution to determine which categories were significantly overrepresented or underrepresented in each group. Significance was determined by Fisher’s Exact test.

### Quantitative RT-PCR

Quantitative reverse transcriptase PCR (qRT-PCR) was conducted to validate the expression of selected genes identified through the RNA-seq analysis. RNA from Arabidopsis leaves and roots from four groups; control, drought stress, nematode infection, and joint stress was treated with DNase I (TURBO DNA-free Kit), and its concentration and purity were determined using a NanoDrop ND-1000 spectrophotometer. cDNA was generated using an iScript cDNA synthesis kit (Bio-Rad).

Primer-Blast (https://www.ncbi.nlm.nih.gov/tools/primer-blast/) was used to design the qRT-PCR primers. The Arabidopsis housekeeping genes *ACTIN2* and *GAPDH* were used for normalisation. Reactions were carried out using SsoAdvanced SYBR green supermix (Bio-Rad). The 2^-ΔΔCt^ method was used to calculate relative expression between samples according to Taylor *et al*. (2019) with three biological and three technical replicates per condition.

### Stress treatments of loss-of-function mutants

The impact of root-knot nematode infection and drought stress either individually or concurrently was investigated for Arabidopsis plants grown in soil. Loss-of-function mutants for the selected candidate genes (*AZI1*, *SAUR71* and *DNR1*) were used for further analysis. T-DNA insertion mutants were obtained from the Nottingham Arabidopsis Stock Centre (NASC). The 35S::*AZI1* homozygous over-expression line was previously generated by Atkinson *et al*. (2013). Seeds of wild type Col-0, mutants, and 35S::*AZI1* were sown on compost and seedlings transplanted into 24-cell trays after two weeks to a mixture of compost:sand:loam at a ratio of 2:1:1. Four weeks old seedlings of each genotype were divided into four groups: Control; Arabidopsis seedlings without any stress treatment, Root-knot nematode infection; Arabidopsis seedlings were infected with 100 J2 *M. incognita* per plant, Drought stress; Water was withheld from the Arabidopsis seedlings until the soil moisture fell to 15-20%, and Joint stress; Arabidopsis seedlings were subjected to combined stress by infection of 100 J2 *M. incognita* per plant and reduction of soil moisture to 15-20%. Ten-twelve plants were used per treatment. A set of growth parameters were measured on different days following treatments. These were rosette diameter, number of rosette leaves, number of rosette branches, height of primary inflorescence, number of siliques per plant.

### Statistical analysis

Statistical analysis was performed using GraphPad Prism (8.2.1). When comparing more than two groups with one independent variable, one-way ANOVA and Tukey post hoc tests were used to determine the significant differences. Two-way ANOVA and Tukey post hoc tests were used if the comparison contained two independent variables. A corrected P value of <0.05 was considered statistically significant. Asterisks indicate significant differences (* P≤ 0.05, ** P ≤ 0.01, *** P≤ 0.001, **** P≤ 0.0001) after the statistical analysis.

## Results

### RNA sequencing of *Arabidopsis thaliana* in response to different stress treatments

To understand plant responses to a combination of drought stress and root-knot nematode infection, *Arabidopsis* plants were either infected with the root knot nematode *M. incognita,* subjected to dehydration or subjected to the two stresses concurrently (joint stress). Total RNA was extracted separately from leaf and root tissues from different treatment groups for further analyses.

Following the assessment of RNA quality, the RNA samples were used for Illumina sequencing. Each sample produced more than 19 million reads and yielded over 5000 million bases. Multidimensional scaling (MDS) plots of RNAseq reads from leaf and root samples confirm that the biological replicates cluster well together and, in all except one instance (control and nematode-infected roots), are separated from other treatments (Figure 1). As may be expected, the lower impact of the nematode infection led to those samples clustering more closely with the controls, whilst the drought stress and joint-stress samples were more distinct. The relatively low impact of the nematode infection alone was amplified to a larger difference once the drought stress was also applied. This trend was continued in roots where there was some overlap between control and nematode-infected samples. Also, there was a higher degree of variance than for leaves between replicate samples of drought-stressed root tissue.

**Figure 1.**
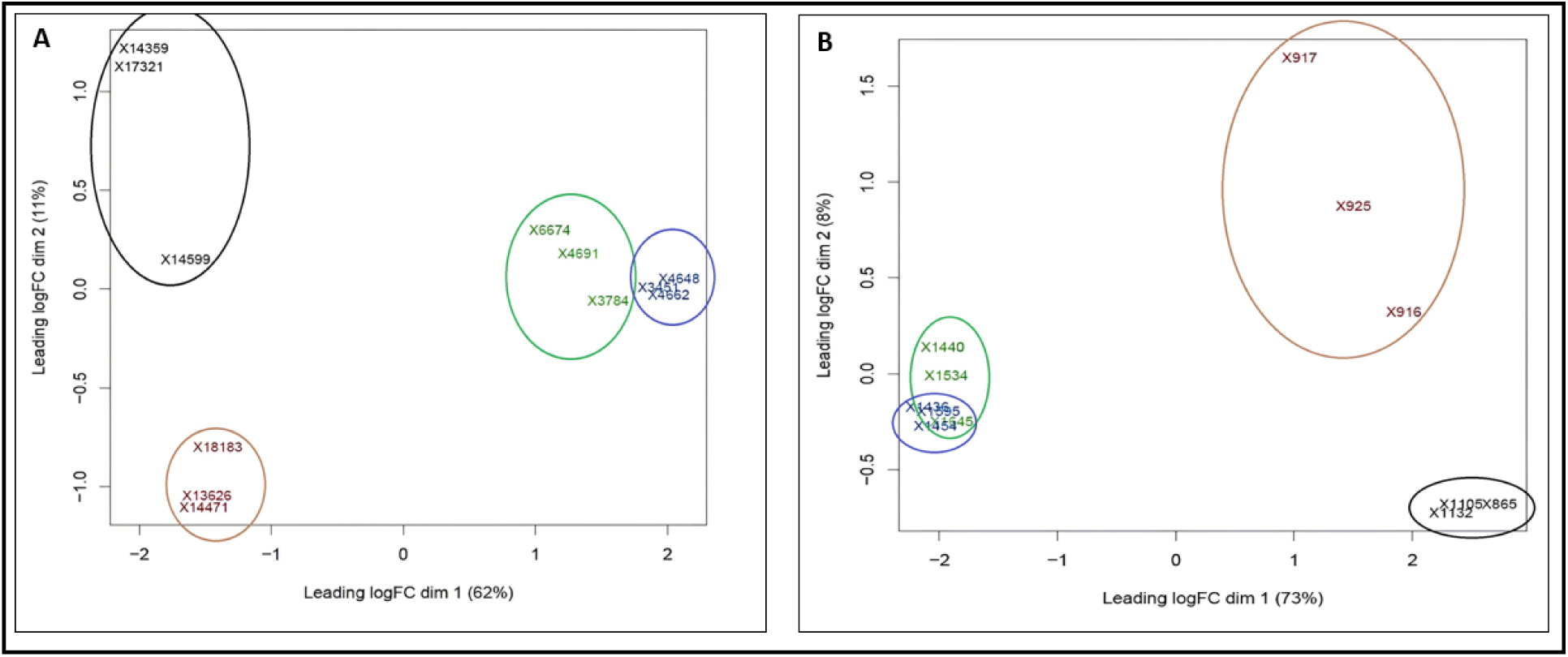
Multidimensional scaling (MDS) plots of sequences from leaf and root samples. Samples within the same treatment group are circled together (Control = blue, nematode infected = green, drought stress = red, and joint stress = black). (A) MDS plot of leaves. (B) MDS plot of roots.

### Identification of differentially expressed genes in Arabidopsis

Differential expression analysis was performed to assess gene expression changes in the leaf and root tissues of Arabidopsis plants subjected to drought stress, root-knot nematode infection and their combination. Genes were classified as differentially expressed if they significantly differed from the control at FDR < 0.05. Table 1 summarises the number of differentially expressed genes (DEGs) for each treatment. In response to nematode infection, 491 genes were differentially expressed in leaves (381 upregulated, 110 downregulated), and 70 in roots (45 up, 25 down). Among root DEGs, approximately 29% of the upregulated genes showed greater than 2-fold induction, whereas none of the downregulated genes reached this threshold. In leaves, 27% of upregulated genes exceeded a 2-fold increase, while only two downregulated genes showed similar magnitude of change.

**Table 1.** Number of differentially expressed genes (DEGs) in leaves and roots in response to different stress treatments. . Genes were included if their expression was significantly different from the control (FDR < 0.05).

| Treatment | Tissue | Number of significantly up-regulated genes | Number of significantly down-regulated genes |
| --- | --- | --- | --- |
| Root-knot nematodes | Leaf | 381 | 110 |
| Drought stress | Leaf | 4901 | 4230 |
| Joint stress | Leaf | 5280 | 4428 |
| Root-knot nematodes | Root | 45 | 25 |
| Drought stress | Root | 7434 | 6933 |
| Joint stress | Root | 8158 | 7512 |

Drought stress elicited a much broader transcriptomic response, with 9,131 DEGs in leaves (4,901 up, 4,230 down) and 14,367 in roots (7,434 up, 6,933 down). Unlike nematode infection, drought induced comparable proportions of up- and downregulated genes in both tissues. A more substantial share of these genes also showed >2-fold changes with 46% of upregulated and 21% of downregulated genes in roots exceeding this threshold. The joint stress treatment triggered a response similar to drought stress alone, though slightly more pronounced. A total of 9,708 genes were differentially expressed in leaves (5,280 up, 4,428 down) and 15,670 in roots (8,158 up, 7,512 down). The distribution of up- and downregulated genes was again balanced, with 46% of upregulated genes in roots showing >2-fold changes. Notably, 82 genes were induced more than 100-fold. Around 27% of downregulated genes also exceeded the 2-fold threshold, including 24 with >100-fold reduction. Those genes with an annotation were classified into different functional categories (biological process, molecular function, cellular component) using The Arabidopsis Information Resource (TAIR) database.

### Overlap between subsets of differentially expressed genes

Venn diagrams illustrate the overlap of differentially expressed genes in Arabidopsis leaves and roots under the different stress treatments (Figure 2). The similarities between Arabidopsis responses to drought stress and nematode infection were lower in roots compared to the leaves. Overall, the transcriptomic responses to drought and nematode infection showed greater similarity in leaves than in roots. In roots, the overlap between DEGs induced by both drought and nematode infection was minimal with fewer than 0.5% of upregulated genes and only 0.2% of downregulated genes shared. In contrast, leaves showed more substantial overlap of responses, with approximately 5.5% of upregulated and 1.7% of downregulated DEGs common to both stresses.

**Figure 2.**
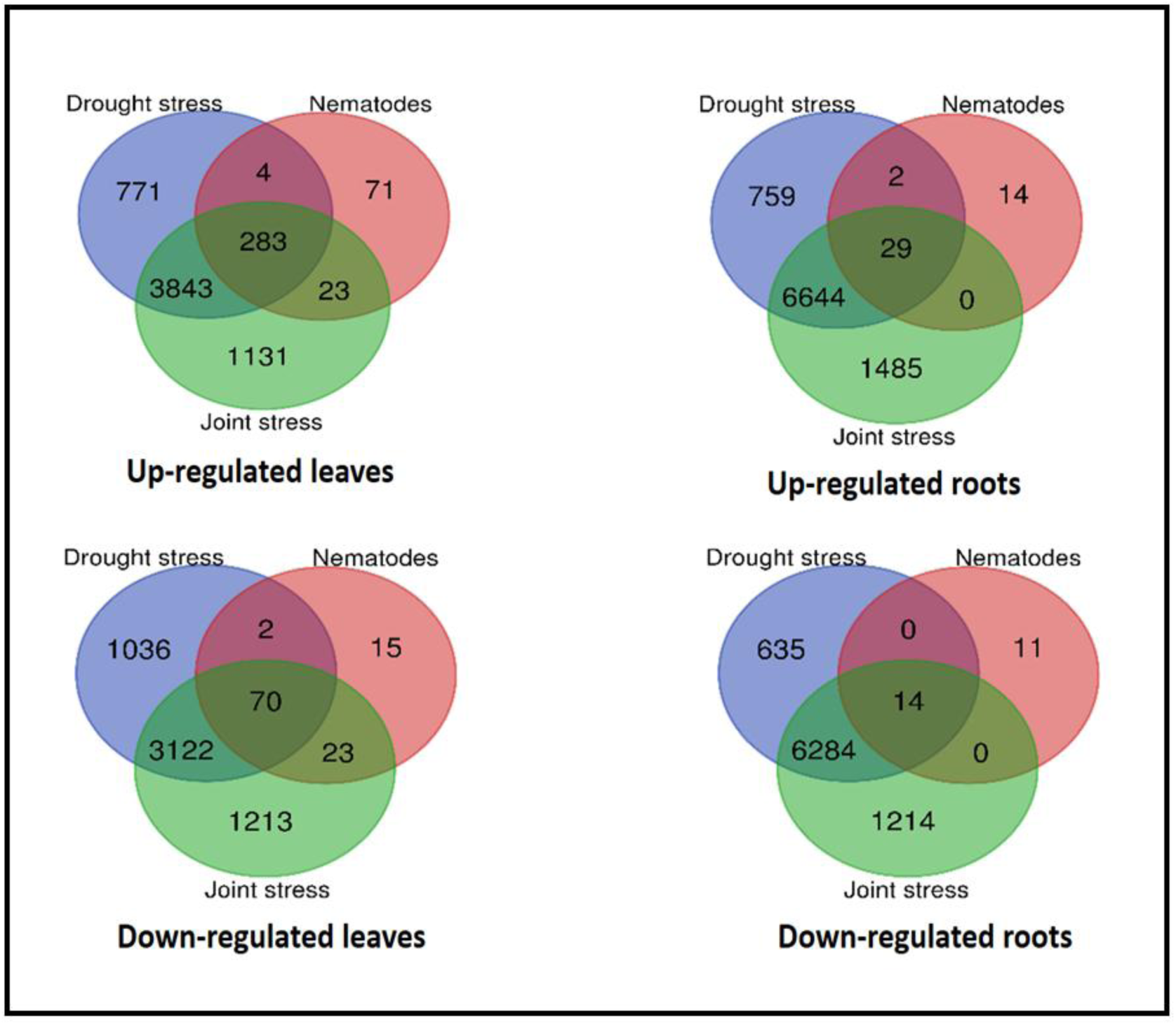
Venn diagrams showing overlap between categories of genes differentially regulated by stress treatments. Genes up- and down-regulated in leaves (A and B, respectively) and roots (C and D, respectively) are shown separately. Genes were included if their expression levels differed significantly from control where FDR < 0.05. Overlapping circles represent genes that were up- or down-regulated by more than one stress treatment.

The combined (joint) stress response shared far more similarity with drought stress than with nematode infection, especially in roots. About 43% of upregulated and 44% of downregulated genes under joint stress were also differentially expressed in response to drought. In comparison, just 0.5% of upregulated and 0.2% of downregulated genes were shared between joint stress and nematode infection. A similar pattern was observed in leaves: 41% of upregulated and 37% of downregulated genes overlapped between drought and joint stress, whereas only 5.4% upregulated and 2.1% downregulated genes overlapped with nematode infection.

In addition to the transcriptomic similarities between the different stress treatments, each stress induced a unique transcriptomic response that was not shared by the other treatments. The unique transcriptomic changes in response to combined drought stress and nematode infection (joint stress) are of particular interest. Despite the overall transcriptomic similarities between joint stress and drought stress, the analysis identified genes that were uniquely regulated in response to the joint stress. There were 1485 genes up-regulated and 1214 genes down-regulated in the roots uniquely by joint stress, whilst 1131 up-regulated and 1213 down-regulated DEGs responded in leaves only to joint stress.

Analysis of the top 50 uniquely up- and downregulated genes under joint stress revealed limited overlap between tissues. Only five genes were commonly upregulated in both leaves and roots (*AT5G24640, AT2G04050, AT3G54530, AT5G54560*, and *AT5G09570*), and just one gene (*AT1G27565*) was commonly downregulated. Gene Ontology (GO) analysis highlighted stress-related functions among these DEGs. In leaves, the ‘defence response’ category included *PLANT DEFENSIN PDF1.1* (AT1G75830), a plant defensin gene upregulated 56.3-fold by joint stress but not by either individual stress. Defensins are antimicrobial peptides with broad-spectrum effects against pathogens. In roots, the ‘response to stress’ category included the highly induced *ARGONAUTE 5 (AGO5)* (AT2G27880). Argonaute family proteins are core components of RNA-induced silencing complexes (RISCs), indicating a potential role of post-transcriptional gene regulation in response to joint stress.

Tables 2 and 3 list the top 10 most highly up- and downregulated unique DEGs in leaves and roots under joint stress. In leaves, *HRE2* (AT2G47520), a hypoxia-responsive ethylene response factor, showed the highest induction (758.2-fold), while the most downregulated gene, encoding a NAD(P)-binding Rossmann-fold superfamily protein (AT1G64590), reduced 33.8-fold. In roots, a gene encoding a beta-galactosidase-related protein was the most highly induced (374.6-fold), and a peroxidase superfamily protein gene (AT1G34510) was the most strongly repressed (149.4-fold).

**Table 2.** The 10 genes that were most highly up-regulated and down-regulated in Arabidopsis leaves, uniquely in response to joint stress. . Genes were included if their expression was significantly different from the control (FDR < 0.05).

| Gene ID | Functional Description | log2FC |
| --- | --- | --- |
| AT2G47520 | Hypoxia responsive ethylene response factor 2, *HRE2* | 9.5665 |
| AT5G54450 | *ATDOB11*, DUF295 ORGANELLAR B 11 | 9.1801 |
| AT3G54530 | Hypothetical protein | 8.9689 |
| AT2G03130 | Ribosomal protein L12/ ATP-dependent Clp protease adaptor protein ClpS family protein | 8.8460 |
| AT5G55150 | F-box SKIP23-like protein (DUF295), *ATFDR2* | 8.7863 |
| AT4G25930 | DUF295 ORGANELLAR A 10, *ATDOA10* | 8.769 |
| AT5G45890 | Senescence-associated gene 12, *SAG12* | 8.143089 |
| AT2G25330 | Tumour necrosis factor receptor-associated factor (TRAF) candidate 1A, TC1A | 7.7324 |
| AT3G54520 | Hypothetical protein | 7.6223 |
| AT5G62480 | Glutathione S-transferase tau 9, *ATGSTU9* | 7.4272 |
| AT1G64590 | NAD(P)-binding Rossmann-fold superfamily protein | -5.11959 |
| AT1G30350 | Pectin lyase-like superfamily protein | -5.01479 |
| AT3G51410 | Hypothetical protein | -5.01215 |
| AT1G27565 | Hypothetical protein | -5.01034 |
| AT5G20330 | *BETAG4*, beta-1,3-glucanase 4 | -4.79105 |
| AT2G19440 | ZERZAUST HOMOLOG, ZETH, O-Glycosyl hydrolase family 17 protein | -4.67012 |
| AT1G72620 | Alpha/beta-Hydrolase superfamily protein | -4.65772 |
| MIR822A | MicroRNA822A, microRNA of unknown function | -3.90653 |
| AT4G09200 | SPLa/Ryanodine receptor (SPRY) domain-containing protein | -3.74215 |
| AT5G35935 | Transposable element gene | -3.66142 |

**Table 3.** The 10 genes that were most highly up-regulated and down-regulated in Arabidopsis roots, uniquely in response to joint stress. . Genes were included if their expression was significantly different from the control (FDR < 0.05).

| Gene ID | Functional Description | log2FC |
| --- | --- | --- |
| AT5G52940 | Hypothetical protein, protein of unknown function (DUF295) | 9.779546 |
| AT4G29200 | Beta-galactosidase related protein | 8.549213 |
| AT5G54560 | Hypothetical protein, Protein of unknown function | 6.713284 |
| AT2G04050 | MATE efflux family protein, | 6.399971 |
| AT2G36800 | DOGT1, don-glucosyltransferase 1 | 6.114344 |
| AT5G26350 | Transposable_element_gene | 5.797552 |
| AT1G71150 | Cyclin-D1-binding protein | 5.733307 |
| AT4G02670 | Indeterminate(ID)-domain 12, IDD12 | 5.702134 |
| AT2G27070 | RR13, response regulator 13, member of Response Regulator: B- Type | 5.596916 |
| AT3G18120 | F-box associated ubiquitination effector family protein | 5.566242 |
| AT1G34510 | PRX8, Peroxidase superfamily protein, Class III peroxidase, | -7.23277 |
| AT1G54970 | PRP1, proline-rich protein 1, encodes a proline-rich protein that is specifically expressed in the root. | -6.48675 |
| AT3G30580 | Hypothetical protein | -6.15245 |
| AT1G54820 | Protein kinase superfamily protein | -5.47031 |
| AT5G57530 | XTH12, xyloglucan endotransglucosylase/hydrolase 12 | -5.40697 |
| AT4G12530 | AZI7, Encodes a member of the AZI family of lipid transfer proteins, Bifunctional inhibitor/lipid-transfer protein/seed storage 2S albumin superfamily protein | -5.39462 |
| AT1G05240 | Pathogenesis related 9, PR9, Peroxidase superfamily protein | -5.38132 |
| AT2G29000 | Leucine-rich repeat protein kinase family protein | -5.36933 |
| AT5G07190 | ATS3, seed gene 3, embryo-specific protein 3 | -5.29602 |
| AT2G23347 | MIR844A, Encodes a microRNA that targets a ubiquitin ligase complex-associated protein kinase family member | -5.08144 |

### Validation of RNA-Seq data by quantitative real–time PCR

Quantitative reverse transcriptase PCR (qRT-PCR) was carried out to verify the differential expression of seven selected genes (*Azelaic Acid Induced 1 (AZI1*, AT4G12470), *Small Auxin Upregulated 71 (SAUR 71*, AT1G56150), *Hypoxia Responsive Universal Stress Protein 1 (HRU1*, AT3G03270), *RGA Target 1 (RGAT1*, AT1G19530), and *Wound-Induced Polypeptide 4 (WIP4*, AT4G10270) in response to the three treatments in both leaves and roots. The expression trends for all genes correlated between the qPCR and RNA-seq analysis (Figure 3).

**Figure 3.**
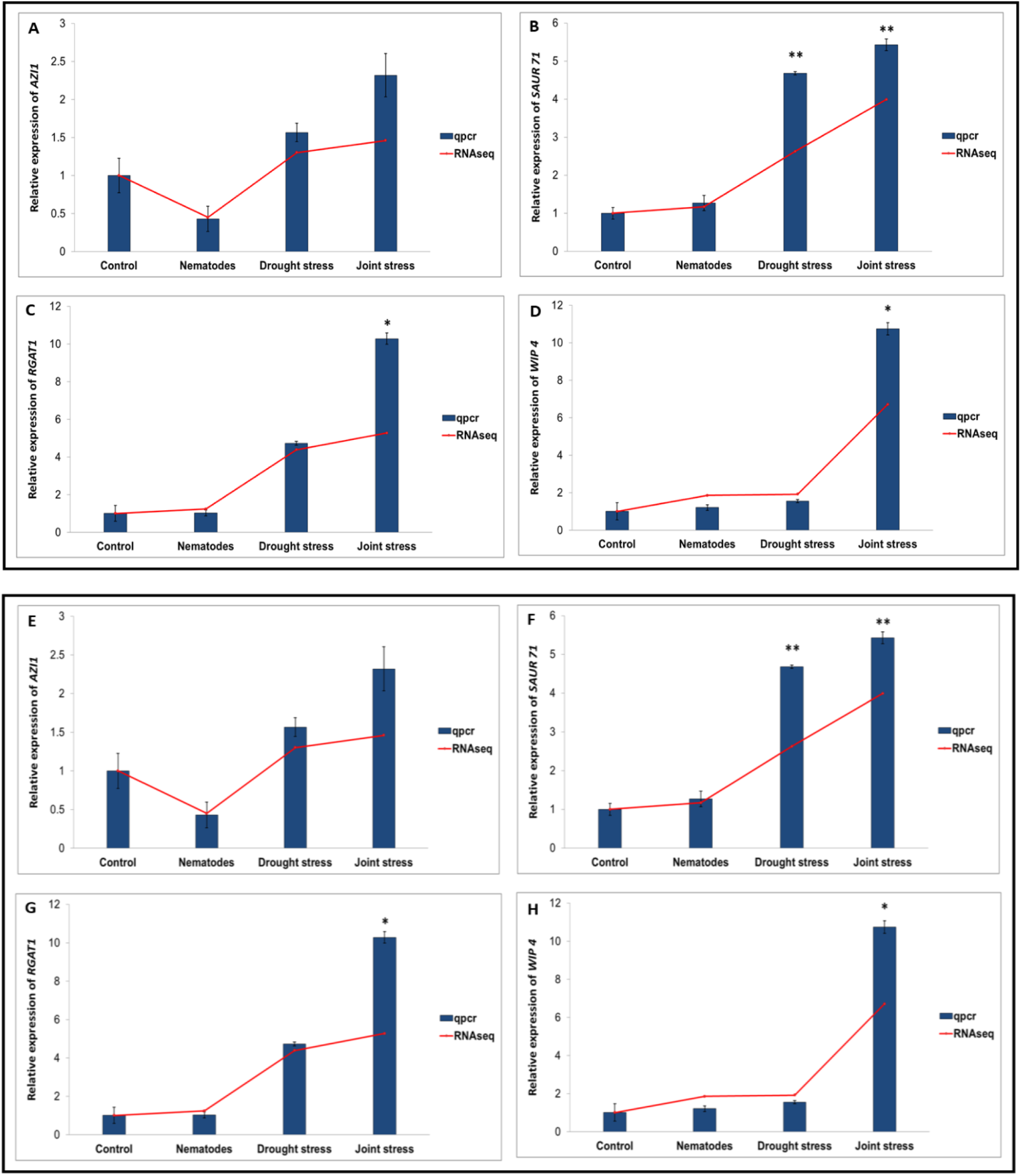
The relative expression of candidate genes in Arabidopsis roots under different stress treatments. The expression levels of (A) *AZI1*, (B) *SAUR71*, (C) *RGAT1,* and (D) *WIP4* Ain leaves. The expression levels of (E) *AZI1*, (F) *SAUR71*, (G) *RGAT1*, and (H) *HRU1* in roots. The expression levels are shown relative to the control, as analyzed by qRT-PCR. Asterisks show a significant difference from the control plants as analysed by one-way ANOVA (*p ≤ 0.05, **p ≤ 0.01). Error bars indicate standard error of the mean. The red line indicates relative expression derived from RNAseq analysis.

### Selection of candidate genes for further analysis

Among the genes uniquely regulated in response to joint stress, *Azelaic Acid Induced1 (AZI1), Small Auxin Upregulated RNA 71 (SAUR 71),* and *Disease Related Nonspecific Lipid Transfer Protein 1 (DRN1)* may play important roles in plant responses to concurrent drought and nematode stresses. *AZI1* and *SAUR71* were both up-regulated in leaves whilst *DRN1* was down-regulated in roots (Table 4). These three candidate genes were selected for further analysis.

**Table 4.** Genes of interest selected from RNA-Seq data for further analysis. Asterisks indicate a significant difference from the control plants (ns =FDR > 0.05, ** =FDR < 0.01). Values less than one represent down-regulation. Values in green boxes represent expression in leaves; values in brown boxes represent expression in roots.

| Gene ID | Gene Name | TAIR Description | Fold change: from control |  |  |
| --- | --- | --- | --- | --- | --- |
|  |  |  | Joint | Drought | RKN |
| AT4G12470 | *Azelaic Acid Induced 1, AZI1* | Induced systemic resistance, systemic acquired resistance | 6.3 ** | 0.7 <sup>ns</sup> | 1.4 <sup>ns</sup> |
| AT1G56150 | *Small Auxin Upregulated 71, SAUR71* | Defence response, hormone-mediated signalling pathway, response to oxidative stress, response to wounding | 3.4 ** | 0.5 <sup>ns</sup> | 0.9 <sup>ns</sup> |
| AT2G45180 | *Disease Related Nonspecific Lipid Transfer Protein 1, DRN1* | Innate immune response activating cell surface receptor signalling pathway, response to salt stress | 0.1 ** | 0.7 <sup>ns</sup> | 0.6 <sup>ns</sup> |

### Expression of *AZI1* in response to different stress treatments

The *Azelaic Acid Induced 1* (*AZI1*) gene was significantly upregulated in *Arabidopsis* leaves under joint stress, whereas neither drought nor nematode infection alone affected its expression. Given the established role of *AZI1* in priming plant defences and mediating systemic acquired resistance (SAR) (Jung *et al*., 2009; Cecchini *et al*., 2015), we further examined the expression profiles of key genes involved in salicylic acid (SA) biosynthesis, along with other members of the *AZI* gene family (Figure 4). Among the *AZI* genes, only *AZI1* was specifically induced in leaves in response to the combined stress. Expression levels of *AZI3*, *AZI5*, and *AZI7* remained unchanged in leaves across all stress treatments. Joint stress uniquely induced the expression of *Isochorismate Synthase 1 (ICS1),* which plays a crucial role in the salicylic acid synthesis pathway, but expression of *ICS2* was not significantly altered. Of the phenylalanine ammonia lyase (*PAL*) genes that are involved in salicylic acid synthesis, only *PAL1* was uniquely upregulated by joint stress. However, *PAL2* and *PAL3* were significantly induced by both drought stress and joint stress while *PAL4* responded to all stress treatments. In root tissues, *PAL4*, *ICS2*, *AZI3*, and *AZI6* were downregulated by both drought and joint stress, whereas *PAL2* and *PAL3* were upregulated under the same conditions. Notably, *ICS1* and *AZI7* were specifically repressed in roots under joint stress but not by either individual stress alone, highlighting a root-specific divergence in SA-related gene regulation in response to combined abiotic and biotic stress.

**Figure 4.**
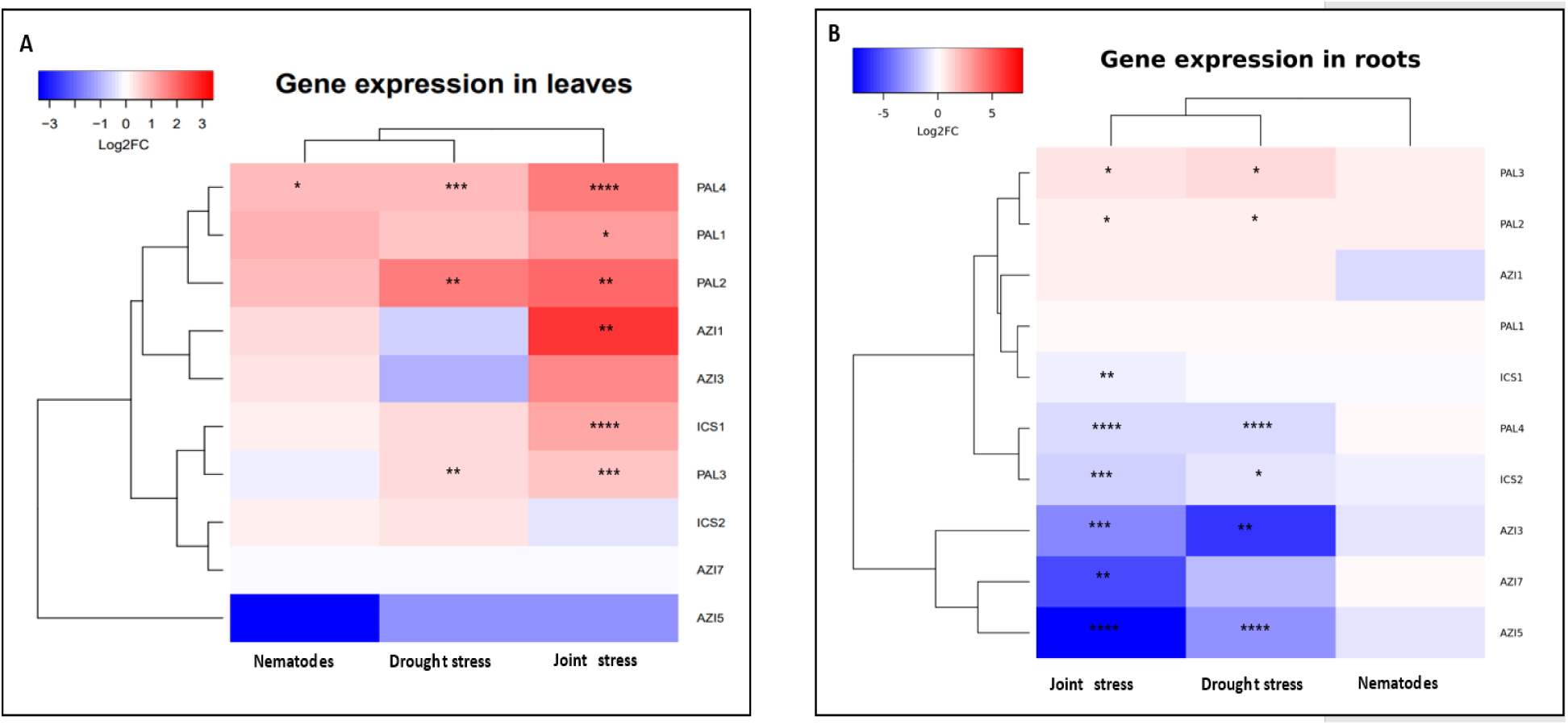
Expression profiles of *AZI* genes and key genes involved in salicylic acid biosynthesis in (A) leaves and (B) roots in response to nematode infection, drought stress, and joint stress relative to control plants based on analysis of RNA-seq data. Asterisks show a significant difference compared to the control plants (*= FDR < 0.05, **= FDR < 0.01, ***= FDR < 0.001, ****= FDR < 0.0001).

### Effect of mannitol and azelaic acid treatments *in vitro*

The effect of drought stress on growth of Arabidopsis WT, *azi1* mutant, and *35S::AZI1* overexpression seedlings was investigated *in vitro*. Six-day old Arabidopsis seedlings were transferred to medium containing 75 mM mannitol for 10 days. Under normal conditions, the fresh weight of *azi1* mutant seedlings after 10 days was significantly higher than WT seedlings (Figure 5A). However, there was no significant difference in primary root length between the two genotypes (Figure 5B). Seedling fresh weight and primary root length were significantly decreased after 10 days exposure to 75 mM mannitol in both WT and *azi1* mutant seedlings (Figure 5A & B). Under mannitol-induced drought stress, the fresh weight of WT and *azi1* plants followed a similar pattern to growth in normal conditions with *azi1* seedlings significantly higher than wild type seedlings. There was no significant difference in primary root length between *azi1* and WT seedlings under mannitol-induced drought stress.

**Figure 5.**
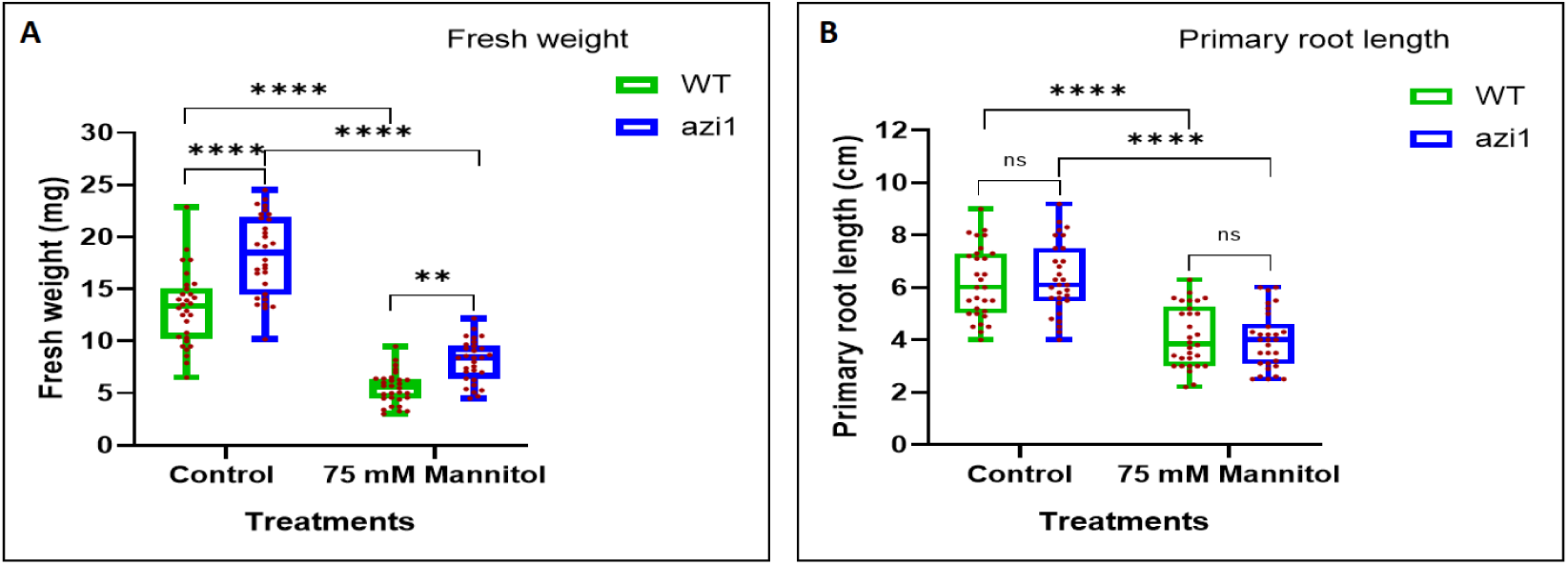
Effect of 75 mM mannitol-induced drought stress on growth parameters of WT Arabidopsis plants and *azi1* mutant line *in vitro*. (A) Fresh weight and (B) Primary root length at 10 days post stress treatments. Asterisks show significant differences (ns > 0.05, **p ≤ 0.01, ****p ≤ 0.0001) as analysed by two-way ANOVA. The significance was determined by the Tukey test (n = 30).

The effect of azelaic acid on Arabidopsis seedlings was investigated *in vitro*. Col-0 Arabidopsis seeds were sown in ½ MS medium supplemented with different concentrations of azelaic acid (10, 30, and 50 µM) or mock for 10 days (Figure 6). Azelaic acid inhibited root growth of Arabidopsis seedlings. Increasing azelaic acid concentrations to 30, and 50 µM significantly decreased the primary root length of Arabidopsis seedlings compared with the mock treatment. There was no significant difference in root growth between low concentration of azelaic acid (10 µM) and the mock treatment.

**Figure 6.**
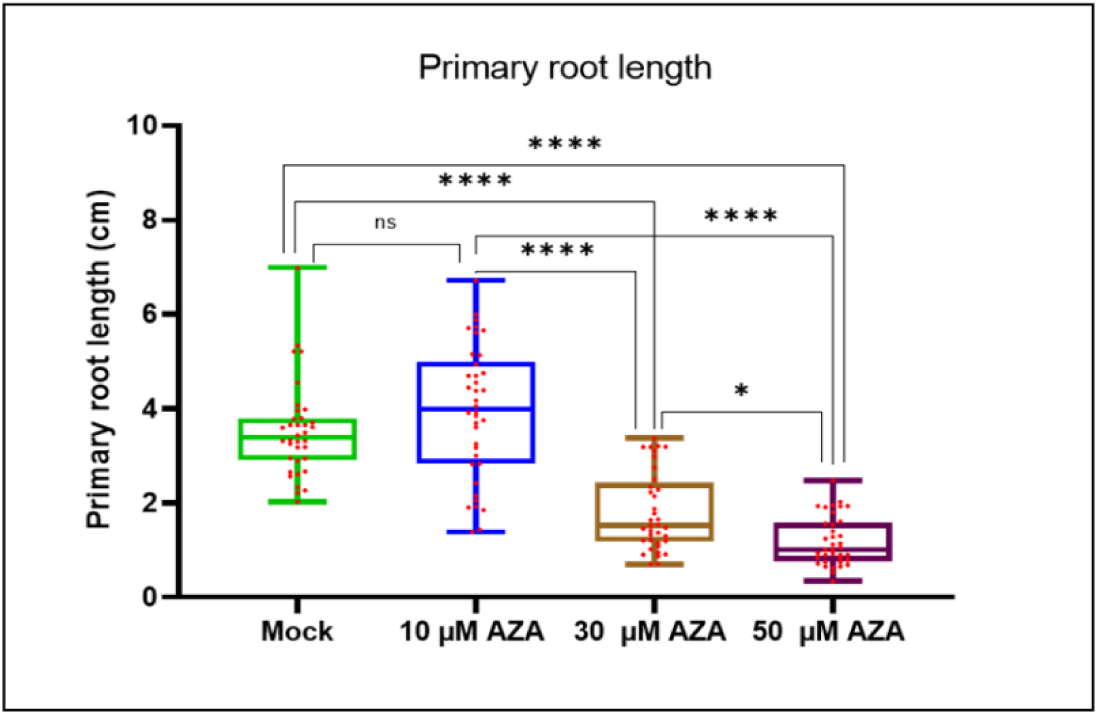
Effect of exogenous Azelaic acid on primary root length of Arabidopsis WT plants grown *in vitro*. Primary root length at 10 days post germination on ½ MS10 medium with and without exogenous azelaic acid. Asterisks show a significant difference from the control plants (ns > 0.05, *p ≤ 0.05, ****p ≤0.0001) as analysed by one-way ANOVA. The significance was determined by the Tukey test (n = 36).

### Expression of the *SAUR* genes in response to different stress treatments

The *SMALL AUXIN UPREGULATED RNA 71* (*SAUR71*) gene was uniquely upregulated in Arabidopsis leaves in response to combined drought stress and root-knot nematode infection. *SAUR71* belongs to the *SAUR* gene family known to play key roles in regulating plant growth and development. To better understand the broader auxin-responsive transcriptional landscape, we extracted expression profiles for all *SAUR* genes in Arabidopsis leaves and roots under individual and combined stress conditions (Figure 7). In leaves, most *SAUR* genes were downregulated by both drought and joint stress, including significant repression of *SAUR1*, *SAUR6*, *SAUR14*, *SAUR16*, *SAUR20*, *SAUR21*, *SAUR50*, *SAUR51*, *SAUR54*, *SAUR62*, *SAUR63*, and *SAUR76*. Notably, nematode infection alone had no significant effect on the expression of these genes. Conversely, several *SAUR* gene (*SAUR34*, *SAUR48*, *SAUR57*, and *SAUR79*) were upregulated in leaves by both drought and joint stress. Strikingly, *SAUR71*, along with *SAUR72* and *SAUR74*, was specifically induced only under joint stress conditions, highlighting a distinct transcriptional response to concurrent biotic and abiotic challenges. Additionally, drought stress alone upregulated *SAUR46* and *SAUR69* and downregulated *SAUR22* in leaves.

**Figure 7.**
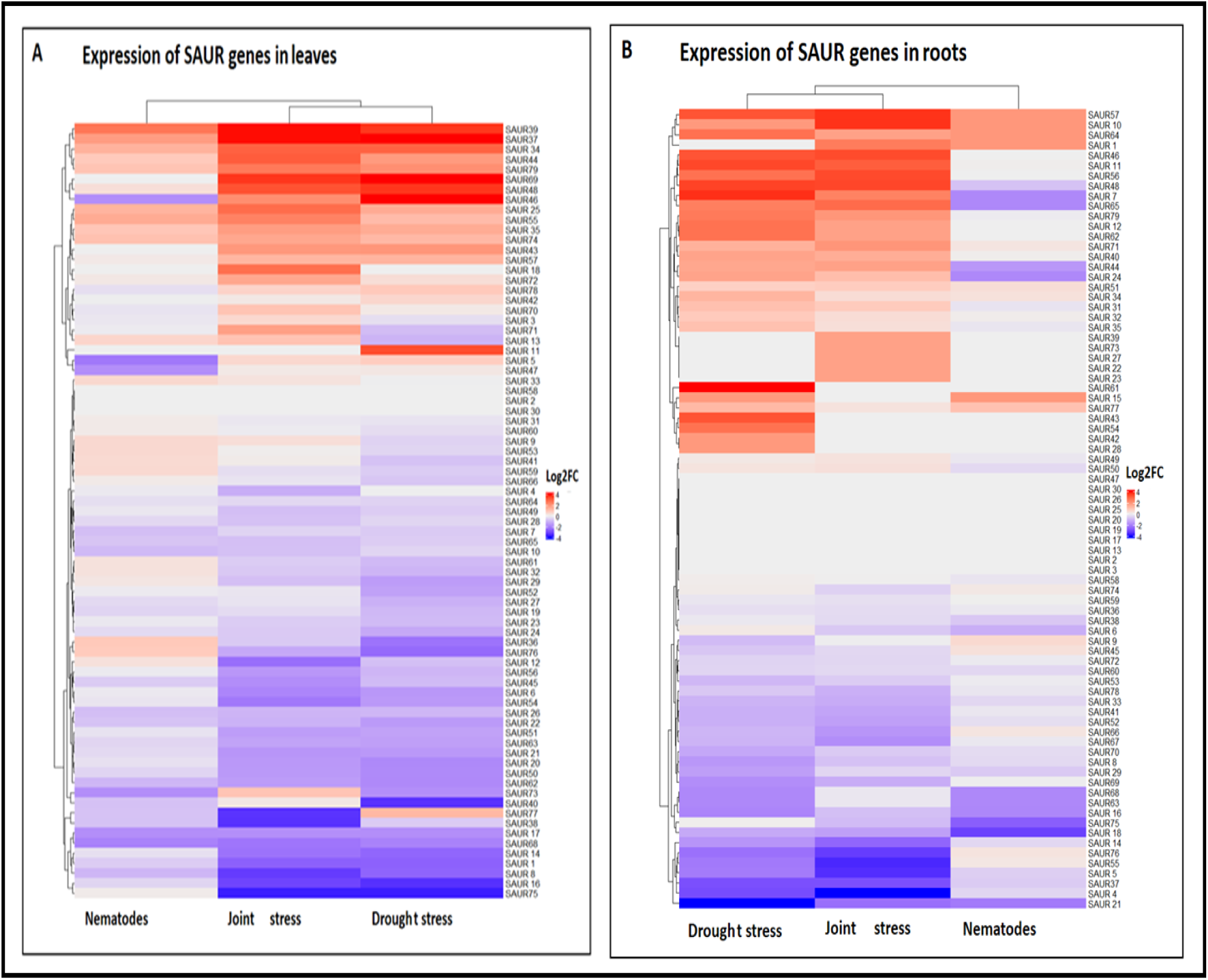
Expression profiles of *SAUR* genes in (A) leaves and (B) roots. The expression of *SAUR* genes in Arabidopsis leaves and roots in response to nematode infection, drought stress, and joint stress relative to control plants based on analysis of RNA-seq data.

In root tissues, fewer *SAUR* genes were affected overall. A subset of *SAUR4*, *SAUR5*, *SAUR33*, *SAUR37*, *SAUR41*, *SAUR52*, *SAUR55*, *SAUR69*, *SAUR70*, *SAUR76*, and *SAUR78* was significantly downregulated under both drought and joint stress. In contrast, *SAUR11*, *SAUR31*, *SAUR32*, *SAUR44*, *SAUR48*, *SAUR71*, and *SAUR79* were significantly upregulated in roots in response to these same treatments. Drought stress alone also induced the expression of *SAUR7*, *SAUR34*, *SAUR35*, and *SAUR77*. Interestingly, *SAUR14* and *SAUR47* were uniquely downregulated in roots under joint stress, further suggesting a specialized response to the combination of environmental pressures.

### Expression of *DRN1* in response to different stress treatments

The *Disease Related Nonspecific Lipid Transfer Protein 1* gene, *DRN1*, was uniquely down-regulated in Arabidopsis roots by joint stress. Its relative expression profile across the three treatments and both tissue types, together with those of other Arabidopsis LTPs, was analysed (Figure 8). Nematode infection alone did not significantly alter the expression of *DRN1* or any other *LTP* genes in either leaves or roots. However, drought and joint stress led to notable changes in *LTP* gene expression, particularly in leaves. Several genes (*LTP1*, *LTP5*, *LTP6*, and *LTP7*) were significantly downregulated by both drought and joint stress. In contrast, *LTP4* was uniquely upregulated in leaves under joint stress. In root tissues, *LTP4* and *LTP5* were downregulated, whereas *LTP2* and *LTP14* were upregulated under both drought and joint stress. The expression of *LTP3* and *LTP10* was down-regulated, while the expression of *LTP8* was up-regulated, only by drought stress in roots. Notably, joint stress uniquely enhanced the expression of *LTP6* while reducing the expression of both *LTP8* and *DRN1* in roots, changes not observed under either stress alone. These findings suggest that specific members of the *LTP* gene family, including *DRN1*, may play targeted roles in *Arabidopsis* responses to combined abiotic and biotic stress.

**Figure 8.**
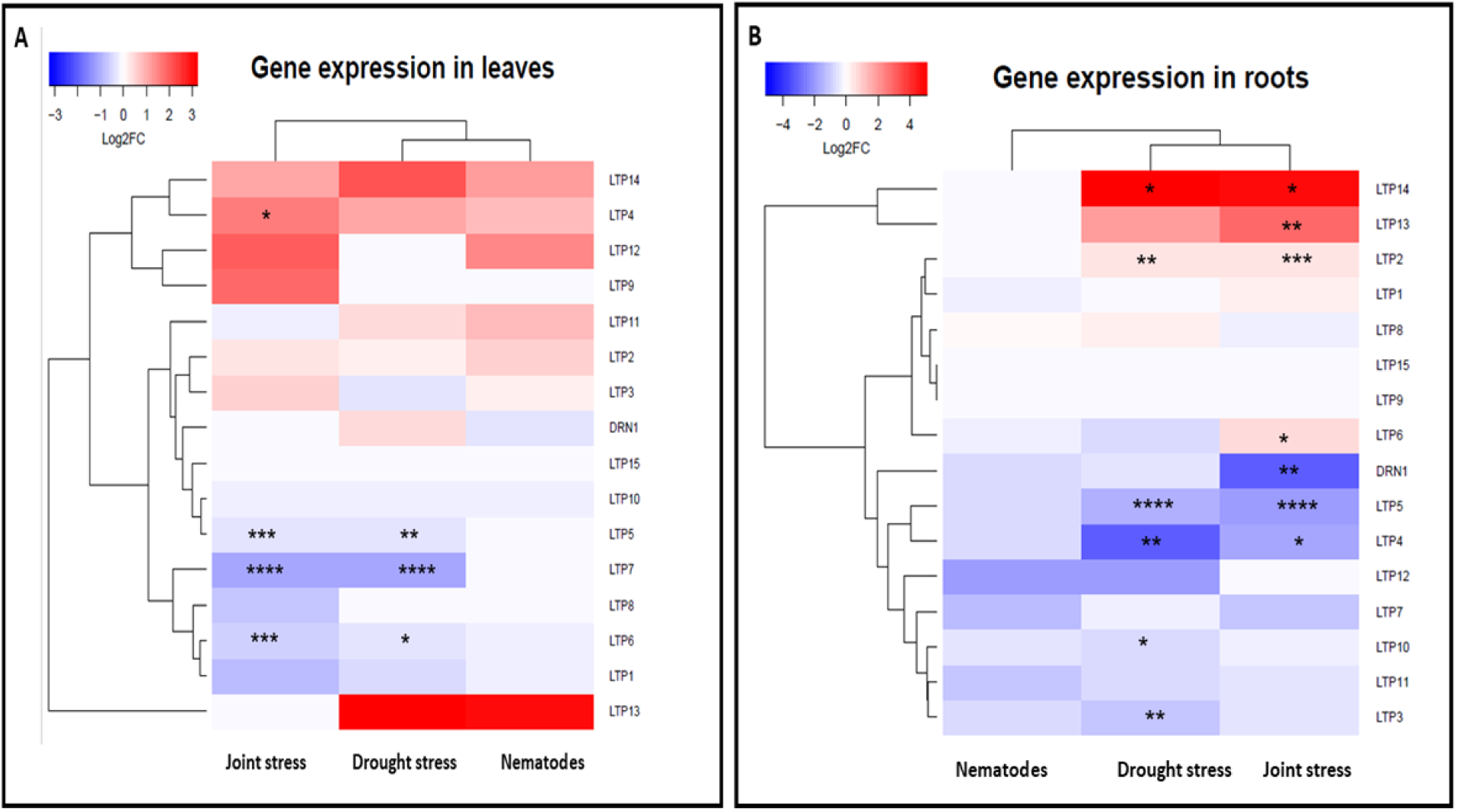
Expression profiles of *DRN1* and other *Lipid Transfer Protein (LTP)* genes in (A) leaves and (B) roots. The expression of *DRN1* and other *LTP* genes in Arabidopsis leaves and roots in response to nematode infection, drought stress and joint stress based on analysis of RNA-seq data. Asterisks show a significant difference relative to the control plants (*FDR < 0.05, **FDR < 0.01, ***FDR < 0.001, ****FDR < 0.0001).

### Response of soil-grown mutant plants to stress treatments

Experiments to investigate the role of AZI1, SAUR71, and DRN1 in plant responses to drought stress and nematode infection either individually or concurrently were carried out using soil-grown wild type and mutant plants. For *AZI1*, a previously characterized over-expression line (Atkinson *et al*., 2013) was also used. Four-week-old seedlings from each genotype were divided into four treatment groups; control, root-knot nematode infection, drought stress, and combined drought stress and nematode infection.

At 8 days after the initiation of stress the rosette diameter of 35S::AZI1 overexpression seedlings in response to joint stress was significantly smaller than for either their control or nematode-infected counterparts (Figure 9A). These differences remained at 16 days following stress treatments, and at this time point joint stress also significantly decreased the rosette diameter of WT plants compared to their control. In contrast, there was no significant impact of any stress treatment on rosette diameter of *azi1* mutants (Figure 9B). At neither time point was there any difference between genotypes exposed to a particular treatment. The number of rosette leaves of each genotype was also measured at 8 days following stress treatments (Figure 9C). Both drought stress and joint stress decreased the rosette leaf number of wild type plants compared to control conditions after 8 days of stress treatments. In addition, the number of rosette leaves of *azi1* mutant plants was significantly decreased under joint stress compared to nematode infection. At the same time point, nematode-infected plants of the 35S::AZI1 overexpression line had fewer rosette leaves than *azi1* mutant plants. The number of rosette branches was measured at 20 days following different stress treatments. The number of rosette branches of WT plants under nematode infection conditions was significantly higher than either control, drought stressed or joint stressed plants. However, here were no significant differences among WT, azi1, and 35S::AZI1 plants under different stress treatments (Figure 9D).

**Figure 9.**
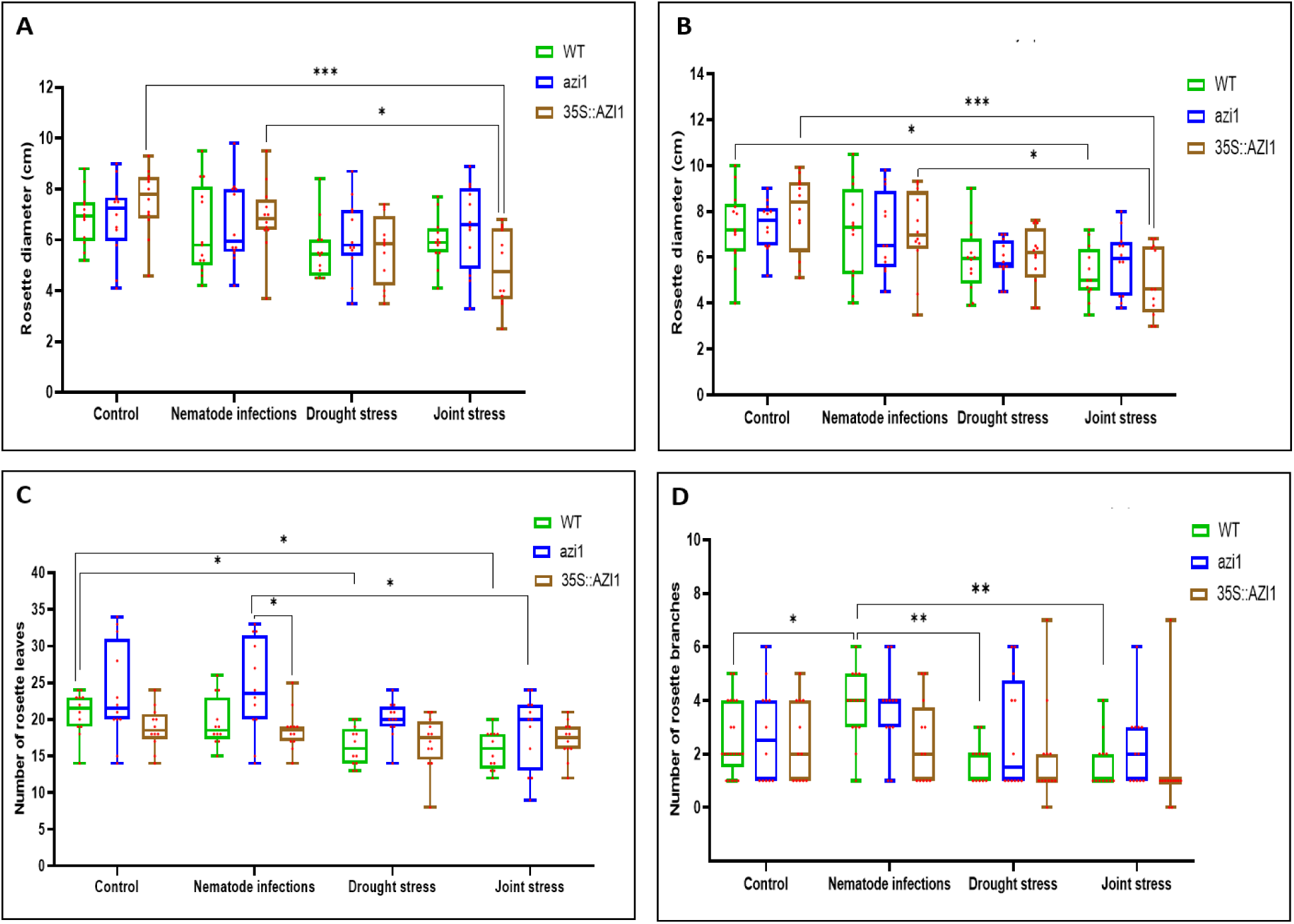
Effect of different stress treatments on growth parameters of Arabidopsis plants WT, azi1 mutant line, and 35S::AZI1 overexpression line. Rosette diameter at (A) 8 days, (B) 16 days post stress treatments. (C) number of rosette leaves at 8 days post stress treatments.(D) number of rosette branches at 20 days post stress treatments. Asterisks show significant differences (*p ≤0.05, **p ≤0.01) as analysed by two-way ANOVA. The significance was determined by the Tukey test (n = 12).

The height of the primary inflorescence of all plants was measured at both 12 and 18 days following initiation of stress treatments. At 12 days, Plants of the 35S::AZI1 overexpression line had consistently shorter inflorescences than the *azi1* mutants in all conditions and were also shorter than WT plants in the absence of drought. In addition, primary inflorescences of the *azi1* mutant plants were significantly taller than those of WT plants whenever nematodes were present. The only treatment-related effect was a significant decrease in height of *azi1* mutants under drought stress compared to nematode infection. By 18 days following initiation of stress treatments, the only within-treatment group difference was consistently shorter primary inflorescences of 35S::AZI1 overexpression plants than *azi1* mutant plants across all conditions. The extent of this was reduced when drought stress was present. Drought stress and joint stress significantly decreased the primary inflorescence height of both WT and *azi1* mutant plants compared to control and nematode-infected conditions (Figure 10 A and B). The total number of siliques per plant was measured at 20 days following different stress treatments. There were no significant differences in silique number between control and different stress treatments for both WT and 35S::AZI1 plants (Figure 10C). Under both control conditions and nematode infection, the number of siliques of *azi1* mutant plants was significantly higher than for 35S::AZI1 overexpression plants. In addition, the number of siliques of *azi1* mutant plants was significantly higher than for WT plants under nematode infection despite that there were no significant differences between them under control conditions. The number of siliques of *azi1* mutant plants under nematode infection conditions was significantly higher than for the control mutant plants and those subjected to drought stress and joint stress.

**Figure 10.**
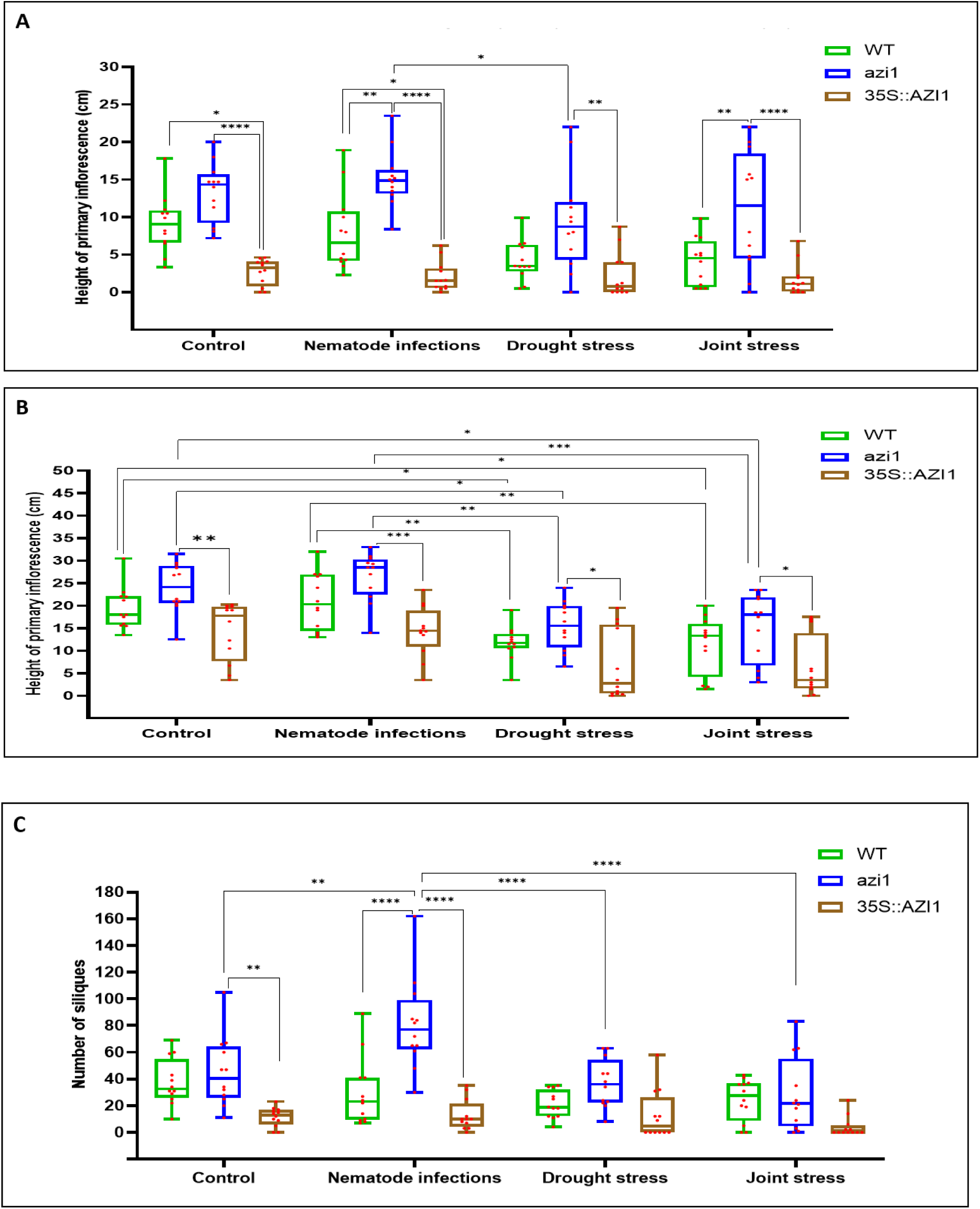
Effect of different stress treatments on growth parameters of WT, *azi1* mutant, and 35S::AZI1 overexpression Arabidopsis plants. Height of primary inflorescence at (A) 12 days and (B) 18 days post stress treatments, and (C) total number of siliques at 20 days post stress treatments. Asterisks show significant differences (*p ≤0.05, **p ≤ 0.01, ***p ≤0.001, ****p ≤0.0001) as analysed by two-way ANOVA. The significance was determined by the Tukey test (n = 12).

For *saur71* mutant plants, there were no significant differences between rosette diameter of wild type and saur71 mutant plants under control and all stress treatments after 8 days (Figure 11A). By 16 days following stress treatments, both drought stress and joint stress treatments significantly decreased the rosette diameter of saur71 mutant plants compared to both control and nematode infection conditions, whilst the wild type plants were impacted only by the joint stress (Figure 11B). There were no significant differences in rosette diameter between wild type and saur71 mutant plants under any conditions. At 8 days following treatments, there was no difference in the number of rosette leaves of wild type and *saur71* mutant plants under any conditions (Figure 11C). Drought stress and joint stress decreased the rosette leaf number of wild type plants compared to control conditions, however only joint stress significantly reduced the number of rosette leaves of the *saur71* mutant plants. There were no significant differences in rosette leaf number between WT and *saur71* mutant plants under control, nematode infection, and joint stress conditions. However, the WT plants had fewer rosette leaves than *saur71* mutant plants under drought stress indicating possible differential susceptibility to drought. The number of rosette branches of *saur71* mutant plants were significantly higher than WT plants under control conditions (Figure 11D). However, there were no significant differences between WT and *saur71* mutant plants under all conditions after 20 days following different stress treatments. Drought stress and joint stress significantly decreased the rosette branches number of *saur71* plants compared to control. Under nematode infections, the number of rosette branches of *saur71* plants was higher than those of drought stress and joint stress treatments. There were no significant differences in primary inflorescence height at 12 days either between WT and saur 71 plants or between the different stress treatments (Figure 12A). By 18 days following stress treatments, both drought stress and joint stress significantly decreased the height of the primary inflorescence of *saur71* plants compared to control plants. The primary inflorescences of nematode-infected *saur71* plants were also significantly taller than those of plants grown under both drought and joint stress conditions (Figure 12B). Both drought stress and joint stress significantly reduced the silique number of *saur71* plants compared to both control and nematode-infected plants (Figure 12C). However, the number of siliques of nematode-infected *saur71* plants was significantly higher than for similarly infected WT plants.

**Figure 11.**
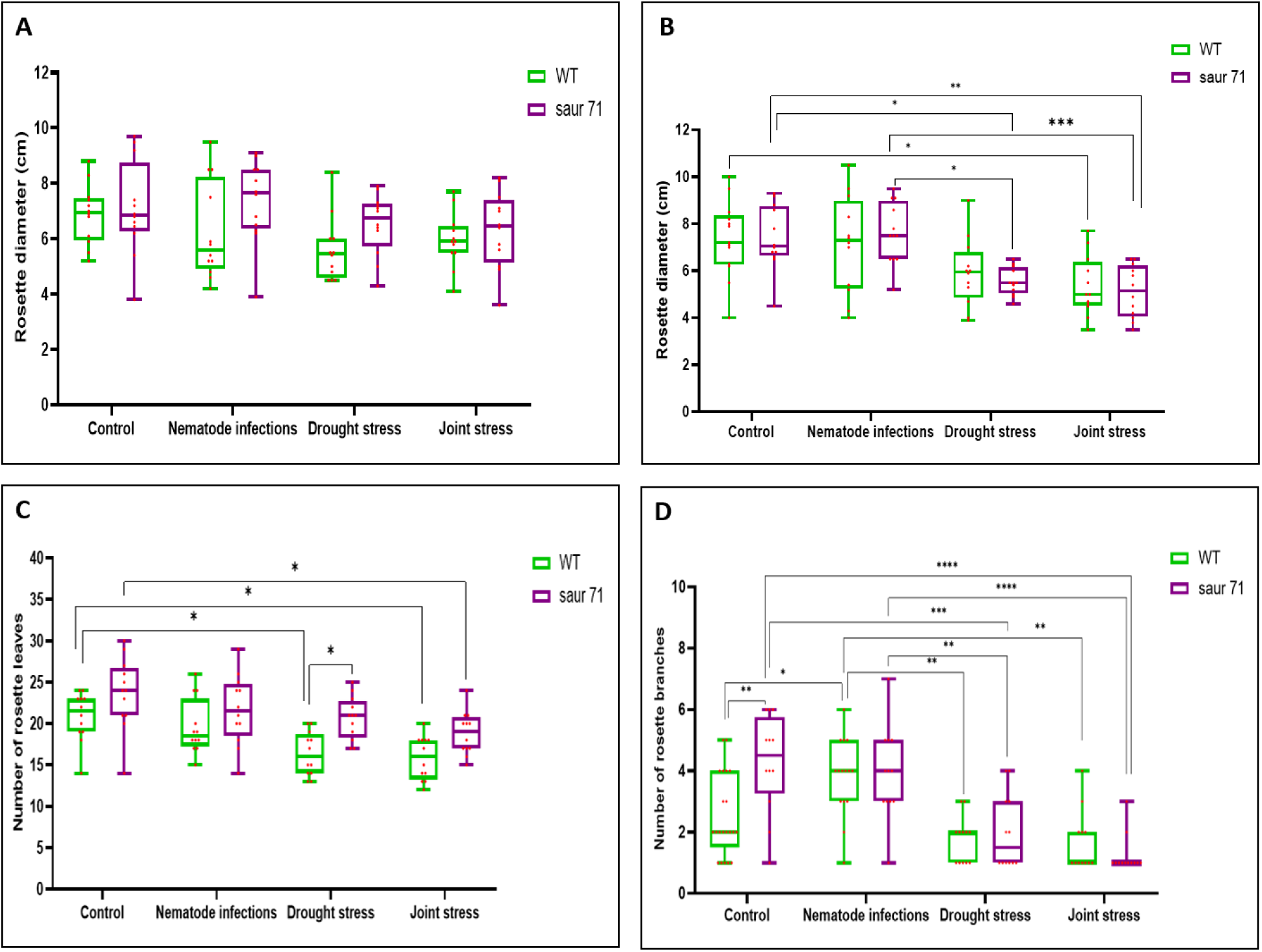
Effect of different stress treatments on growth parameters of WT and *saur 71* mutant Arabidopsis plants. Rosette diameter at (A) 8 days, and (B) 16 days post stress treatments. (C) number of rosette leaves at 8 days post stress treatments. (D) Number of rosette branches at 20 days post stress treatments. Asterisks show significant differences (*p ≤ 0.05, **p ≤ 0.01) as analysed by two-way ANOVA. The significance was determined by the Tukey test (n = 12).

**Figure 12.**
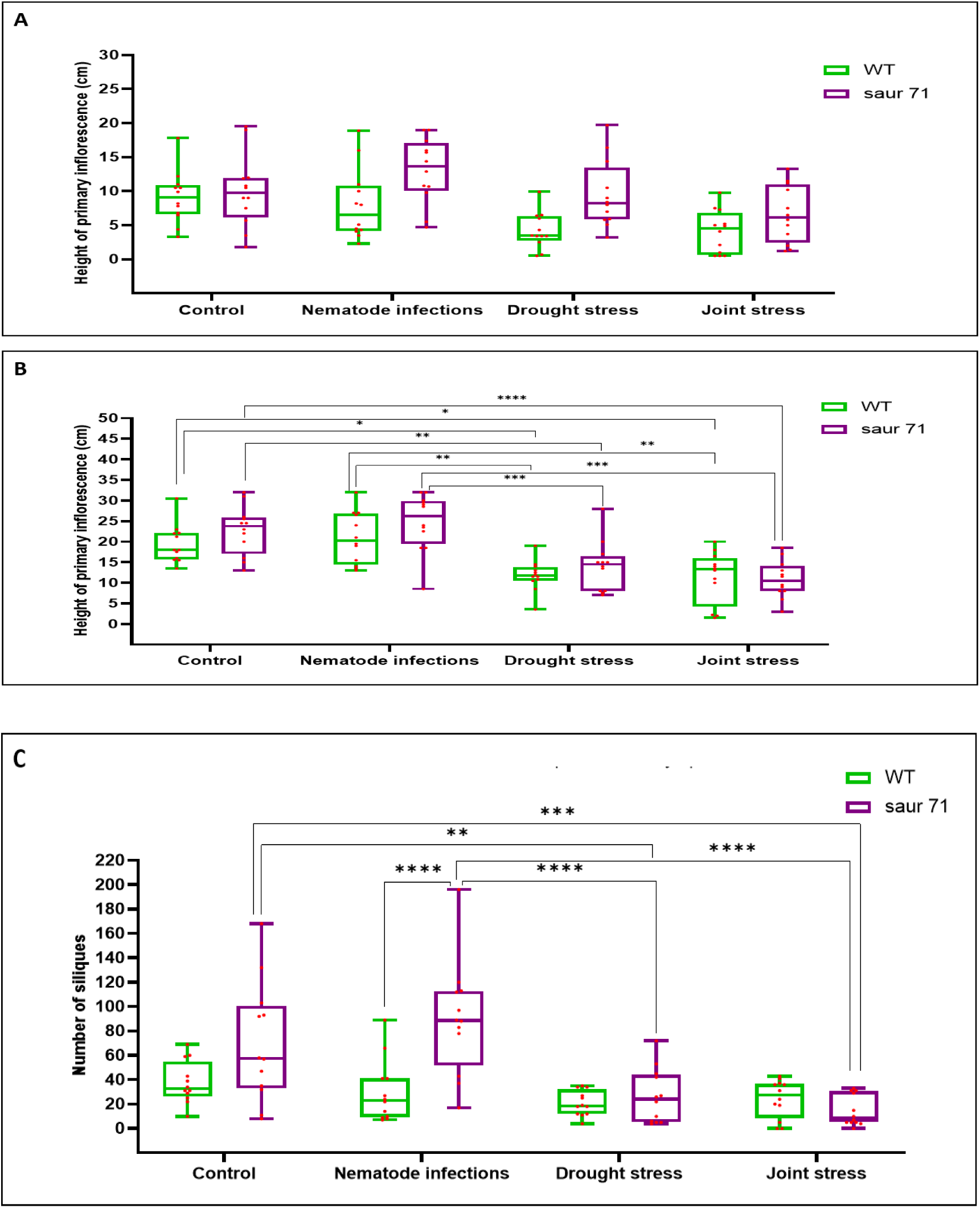
Effect of different stress treatments on growth parameters of WT and *saur 71* mutant Arabidopsis plants. Height of primary inflorescence at (A) 12 days and (B) 18 days post stress treatments, and (C) number of siliques at 20 days post stress treatments. Asterisks show significant differences (*p≤ 0.05, **p≤ 0.01, ***p ≤ 0.001, ****p ≤ 0.0001) as analysed by two-way ANOVA. The significance was determined by the Tukey test (n = 12).

For *drn1* mutant plants, joint stress significantly reduced the rosette diameters of drn1 mutant seedlings compared to control plants after 8 days of treatments (Figure 13A). However, after 16 days only the wild type plants showed a reduced rosette diameter in response to the joint stress(Figure 13B). Drought stress and joint stress decreased the number of rosette leaves of *drn1* plants compared to control conditions after 8 days of stress treatments (Figure 13C). There were no significant differences in the number of rosette branchs between the control and the other stress treatments or among the different stress treatments. In addition to, there were no significant changes in rosette branch number between WT and *drn1* mutant plants under control, nematode infection, drought stress and joint stress conditions (Figure 13D). Joint stress significantly decreased the primary inflorescence height of *drn1* mutant plants compared to both control and nematode infected plants. However, there were no significant differences between WT and *drn1* mutant plants under control and all stress treatments Figure 14A). By 18 days of stress treatments, both drought stress and joint stress decreased the height of primary inflorescence of *drn1* plants compared to control conditions (Figure 14B). In addition to, the number of siliques of *drn1* plants was significantly decreased under both drought stress and joint stress conditions compared to control plants (Figure 14C). In contrast, there were no significant changes in the siliques number of wild type plants under nematode infection, drought stress or joint stress.

**Figure 13.**
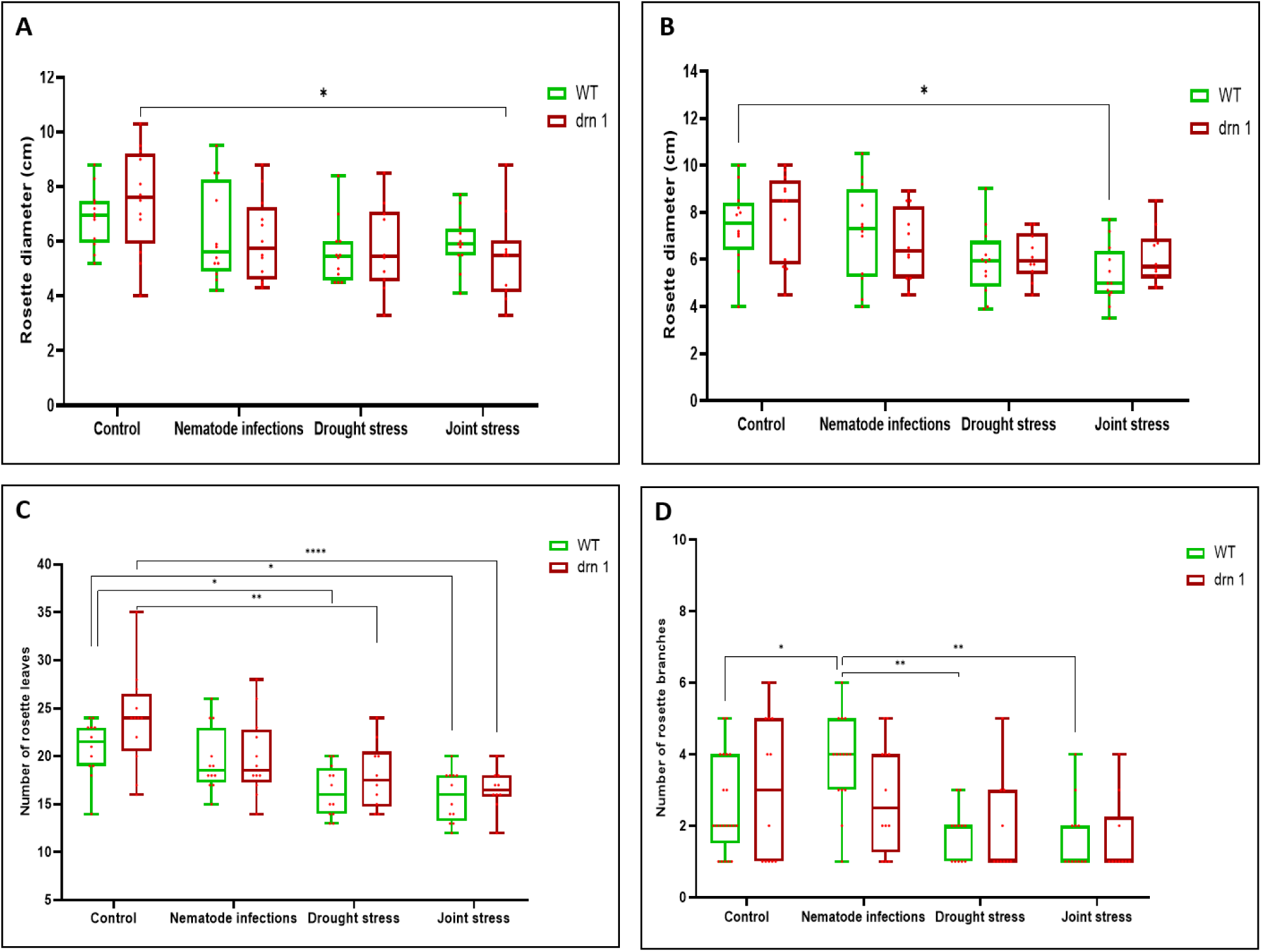
Effect of different stress treatments on growth parameters of WT and *drn1* mutant Arabidopsis plants. Rosette diameter at (A) 8 days, and (B) 16 days post stress treatments. (C) Number of rosette leaves at 8 days post stress treatments. (D) Number of rosette branches at 20 days post stress treatments. Asterisks show significant differences (*p ≤0.05, **p ≤ 0.01, ****p ≤ 0.0001) as analysed by two-way ANOVA. The significance was determined by the Tukey test (n = 10-12).

**Figure 14.**
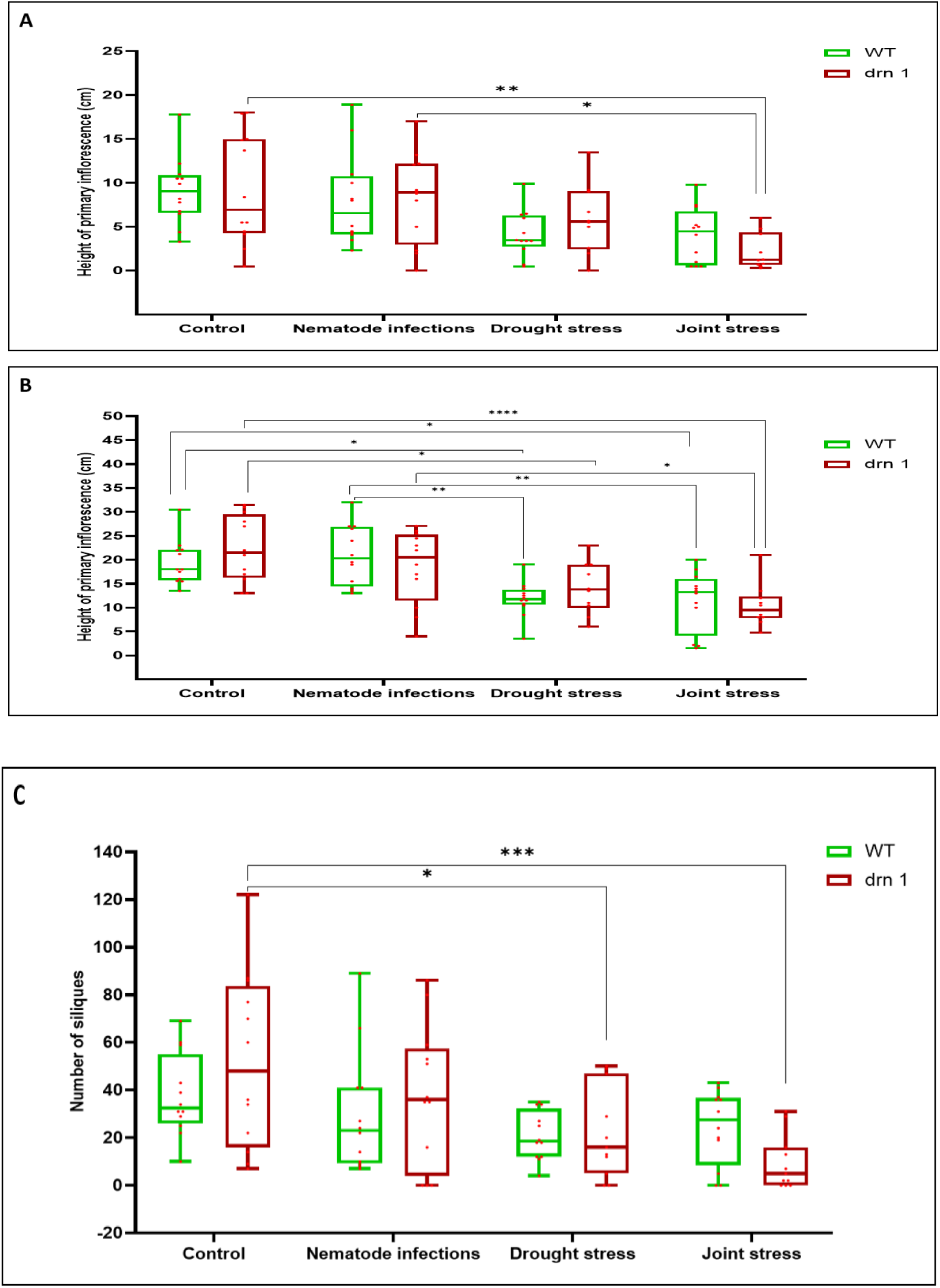
Effect of different stress treatments on growth parameters of Arabidopsis WT and *drn1* mutant plants. Height of primary inflorescence at (A) 12 days and (B) 18 days post stress treatments, and (C) number of siliques at 20 days post stress treatments. Asterisks show significant differences (*p≤ 0.05, **p≤ 0.01, ***p ≤ 0.001, ****p ≤ 0.0001) as analysed by two-way ANOVA. The significance was determined by the Tukey test (n = 10-12).

## Discussion

RNA-seq analysis is widely used as a powerful technique to understand the molecular changes in plants in response to different biotic and abiotic stress factors. Transcriptome analysis has been instrumental in understanding plant-pathogen interactions and investigating the molecular responses of plants to a wide range biotic and abiotic stress factors. In this study, RNA-seq analysis was used to investigate the transcriptomic changes in Arabidopsis plants grown *in vitro* in response to root-knot nematodes and drought stress, either individually or concurrently. The resulting dataset allowed identification of candidate genes that may play a role in plant responses specifically to the combinatorial stress.

### Plant responses to combined stress are complex and tissue-specific

Root-knot nematode infection affected the expression of 70 genes in roots and 491 in leaves, corresponding to ∼0.2% and ∼1.5% of annotated Arabidopsis genes, respectively. Interestingly, nematode-induced transcriptomic changes were more pronounced in leaves than roots, highlighting stronger systemic responses in distant tissues compared to local responses. This contrasts with a previous microarray study of cyst nematodes (*H. schachtii*) infecting Arabidopsis *in vitro*, where more balanced expression changes across leaves and roots were reported. At 10 dpi, 19 genes were upregulated and 28 downregulated in roots, compared to 56 upregulated and 28 downregulated in leaves (Atkinson *et al*., 2013). The number of genes with altered expression in that study was also lower than reported here using the same experimental set-up, likely a consequence of either the different infecting nematode species, or the difference in sensitivity between microarray and RNAseq approaches. Both studies analysed whole root systems, which may mask localised gene expression patterns. In contrast, transcriptomic analyses focused on root-knot nematode-induced galls reveal significantly higher and distinct DEG profiles. For instance, *M. incognita*-induced galls in Arabidopsis contained 1249, 4161, and 4927 DEGs at 3, 5, and 7 dpi, respectively (Yamaguchi *et al.,* 2017) whilst similar transcriptomic patterns were reported by Jammes *et al*. (2005). However, because galls represent a small portion of the root encompassing the feeding site, they do not fully capture systemic responses. Thus, whole-root analyses, as used in this study, provide a more representative overview of the broader molecular changes induced by nematodes, whilst also enabling accurate comparisons across stress treatments.

### Transcriptomic responses to drought and combined stresses

Drought stress, simulated in this work by air-drying *in vitro*, induced widespread gene expression changes: 14,367 DEGs in roots (∼42% of annotated genes) and 9131 in leaves (∼27%). This extensive response, particularly in roots, aligns with their role as primary drought sensors. The relatively short and intense drought treatment likely caused early root responses to predominate, with the response in leaves reflecting the fact that roots generate long-distance signals to initiate systemic molecular drought responses (Janiak et al., 2016; Rasheed et al., 2016). Combined drought and nematode stress triggered 15,670 DEGs in roots (∼46%) and 9708 in leaves (∼29%). However, only a small fraction of these overlapped with the nematode-only response - just 43 DEGs (0.27%) in roots and 399 DEGs (4.1%) in leaves. In contrast, drought was the dominant driver of transcriptomic changes under combined stress, with 12,971 DEGs in roots (82.8%) and 7318 in leaves (75.4%) overlapping with the drought-only response. These findings suggest that under combined stress conditions, drought has a far stronger influence on global gene expression patterns in both roots and leaves, whereas nematode-specific responses are largely diluted or overridden.

### Combined drought and nematode stress triggers a unique transcriptomic response

Simultaneous drought stress and root-knot nematode infection activated a distinct gene expression program in Arabidopsis that differed substantially from the transcriptomic profiles triggered by either stress alone. This joint stress response was not merely additive, but instead represented a unique molecular signature. Specifically, 2344 DEGs in leaves (∼24.1%) and 2699 DEGs in roots (∼17.2%) were exclusively regulated by the combined stress treatment, not by individual drought or nematode stress. These findings align with previous studies demonstrating that combined abiotic and biotic stresses often induce distinct transcriptomic responses, reflecting the plant’s ability to integrate multiple environmental cues into a specific and targeted response (Rizhsky *et al*., 2002; Atkinson *et al*., 2013; Rivero *et al*., 2014; Kim *et al*., 2022). Among the uniquely regulated genes, *AZI1*, *SAUR71*, and *DRN1* were selected for further investigation. *AZI1*, a key regulator of systemic acquired resistance (SAR), was specifically upregulated in leaves under joint stress. Interestingly, this gene was also differentially expressed in Arabidopsis leaves specifically in response to the joint stress of drought and cyst nematode infection, however in that case *AZI1* was down-regulated (Atkinson *et al*., 2013). *SAUR71*, a member of the largest family of primary auxin-response genes, also showed unique induction. In contrast, *DRN1*, a member of the non-specific lipid-transfer protein family, was strongly downregulated in roots. These gene expression patterns suggest the activation of novel defence and signalling pathways in response to the combined stress, that may be specific to root-knot nematode infection.

### *AZI1* and the salicylic acid pathway in joint stress response

AZI1 is a crucial mediator of defence priming and SAR (Jung *et al*., 2009; Cecchini *et al*., 2015), processes that enhance a plant’s ability to mount rapid and robust responses to stress. A primed plant is in a physiological state whereby it can respond more quickly and strongly to future stress. Plant performance is improved, and resistance and tolerance are enhanced without fully activating defence pathways until the plant is challenged again (Conrath *et al*., 2006; Mauch-Mani *et al*., 2017). Induction of *AZI* is driven by azelaic acid, a mobile metabolite transported via plasmodesmata (Lim *et al.,* 2016) that promotes SAR (Parker, 2009) and elevates levels of salicylic acid (SA) (Jung *et al.,* 2009) a central hormone in biotic stress responses.

The starting point of SA biosynthesis is chorismate, which is converted to either isochorismate through the isochorismate synthase (ICS) pathway or to prephenate via the phenylalanine ammonia lyase (PAL) pathway (Lefevre *et al.,* 2020; van Butselaar & van den Ackerveken, 2020). To explore the link between *AZI1* and SA biosynthesis, expression of *isochorismate synthase 1* (*ICS1*) and phenylalanine ammonia lyase (*PAL*) genes was analysed. *ICS1*, a key enzyme in the ICS pathway of SA biosynthesis, was uniquely upregulated by joint stress. Similarly, *PAL1* showed specific induction by joint stress while other *PAL* genes exhibited varied responses: *PAL4* was induced by all treatments, *PAL2* and *PAL3* responded to drought and joint stress, but not nematodes alone. These results suggest that the ICS pathway, rather than the PAL pathway, predominantly contributes to SA production under combined drought and root-knot nematode stress, consistent with its established role in pathogen-induced SAR (Wildermuth *et al.,* 2001; Loake & Grant, 2007). SA is essential for defence against biotrophic pathogens including root-knot nematodes. For example, *Mi-1*-mediated resistance in tomato against *M. javanica* is SA-dependent, and loss of SA biosynthesis impairs SAR and pathogen resistance (Branch *et al.,* 2004).

*AZI1* expression and azelaic acid signalling are also linked to growth suppression. Arabidopsis plants with loss-of-function mutations in *AZI1* showed enhanced growth, while exogenous azelaic acid inhibited root growth, highlighting a growth-defence trade-off. This mechanism may serve to limit the number of nematode feeding sites and reduce water loss, thus improving stress tolerance. In this study, azelaic acid-induced defence prioritisation was specific to the combined stress condition and not observed with individual treatments. Plants must balance growth and defence due to limited resources, and this trade-off is regulated by complex hormonal crosstalk. Growth-promoting and defence-related hormones interact, often antagonistically, to determine physiological outcomes based on the type and severity of stress (Vos *et al*., 2013; Huot *et al*., 2014; van Butselaar & van den Ackerveken, 2020). Under combined drought and nematode stress, this balance appears to shift in favour of defence, as seen by the upregulation of defence-related genes and suppression of growth-associated responses.

### *SAUR* and *DRN1* gene regulation under combined stress

Several *small auxin upregulated RNA* (*SAUR*) genes had altered expression in Arabidopsis under drought, nematode infection, or both. Members of the *SAUR* gene family, the largest group of primary auxin response genes (Stortenbeker & Bemer, 2019), are rapidly induced by auxin and promote cell expansion via activation of plasma membrane (PM) H⁺-ATPases (Spartz *et al*., 2014; Nagpal *et al*., 2022). These enzymes acidify the cell wall by suppressing PP2C-D phosphatases, facilitating growth and potentially influencing plant–microbe interactions as PM H⁺-ATPases are involved in plant immune responses and regulation of stomatal apertures during pathogen infection (Sondergaard *et al.,* 2004; Liu *et al*., 2009; Elmore & Coaker, 2011). Three *SAUR* genes were uniquely upregulated in leaves in response to combined stress, with *SAUR71* showing the highest fold change. Mutant *saur71* plants showed reduced growth under drought and combined stress, but not nematode infection alone. Previous studies support SAURs as key players in stress responses and development, but the situation is complex. For example, overexpression of *SAUR56/60* increased stomatal aperture and promoted stomatal opening, however this suggests that their over-expression would reduce drought tolerance (Wong *et al*., 2021). In contrast over-expression of *SAUR32* improved drought survival via ABA signalling (He *et al.,* 2021). Notably, *SAUR71* is induced by abscisic acid (ABA) rather than auxin (Qiu *et al*., 2020), aligning it with ABA-dependent drought signalling pathways (Ilyas *et al*., 2021; Muhammad Aslam *et al*., 2022).

In contrast, *DRN1*, a gene encoding a non-specific lipid transfer protein (nsLTP), was uniquely downregulated in roots under combined stress. NsLTPs are implicated in membrane stability, signalling, and resistance to a range of stresses (García-Olmedo *et al*., 1995; Missaoui *et al*., 2022). The antimicrobial activity of some nsLTPs supports additional roles in defence against pathogen infections (Kirubakaran *et al*., 2008; Liu *et al*., 2015). *DRN1* suppression has been linked to increased susceptibility to bacterial and fungal infections, and to salt stress (Dhar *et al*., 2020). Its expression is also reduced by reactive oxygen species (ROS), a key signal in drought stress responses, and by salt stress (Obermeyer *et al.,* 2022). In this study, *drn1* mutant plants exhibited reduced growth under combined stress, suggesting DRN1 plays a role in adaptation to overlapping biotic and abiotic stressors.

## Conclusion

Plants in natural environments are frequently exposed to simultaneous biotic and abiotic stresses. This study demonstrates that combined drought stress and root-knot nematode infection elicit unique transcriptomic responses in Arabidopsis, distinct from those triggered by either stress alone. The modulation of specific genes like *AZI1*, *SAUR71*, and *DRN1* under joint stress highlights the complexity and specificity of plant stress responses. These findings reinforce the need to study plant responses under realistic, combined stress conditions to inform breeding strategies aimed at developing resilient, multi-stress-tolerant crops.

## References

Al-Sadi, A.M., Al-Masoudi, R.S., Al-Habsi, N., Al-Said, F.A., Al-Rawahy, S.A., Ahmed, M. and Deadman, M.L. 2010. Effect of salinity on pythium damping-off of cucumber and on the tolerance of Pythium aphanidermatum. Plant Pathology. 59(1), pp.112–120.

Ameye, M., Wertin, T.M., Bauweraerts, I., McGuire, M.A., Teskey, R.O. and Steppe, K. 2012. The effect of induced heat waves on P inus taeda and Q uercus rubra seedlings in ambient and elevated CO2 atmospheres. New Phytol. 196(2), pp.448–461.

Atkinson, N.J., Dew, T.P., Orfila, C. and Urwin, P.E. 2011. Influence of combined biotic and abiotic stress on nutritional quality parameters in tomato (Solanum lycopersicum). J Agric Food Chem. 59(17), pp.9673–9682.

Atkinson, N.J. and Urwin, P.E. 2012. The interaction of plant biotic and abiotic stresses: from genes to the field. J Exp Bot. 63(10), pp.3523–3543.

Atkinson, N.J., Lilley, C.J. and Urwin, P.E. 2013. Identification of genes involved in the response of Arabidopsis to simultaneous biotic and abiotic stresses. Plant Physiol. 162(4), pp.2028–2041.

Audebert, A., Coyne, D.L., Dingkuhn, M. and Plowright, R.A. 2000. The influence of cyst nematodes (Heterodera sacchari) and drought on water relations and growth of upland rice in Cote d’Ivoire. Plant Soil. 220(1-2), pp.235–242.

Bidzinski, P., Ballini, E., Ducasse, A., Michel, C., Zuluaga, P., Genga, A., Chiozzotto, R. and Morel, J.B. 2016. Transcriptional basis of drought-induced susceptibility to the rice blast fungus Magnaporthe oryzae. Front Plant Sci. 7, p.1558.

Branch, C., Hwang, C.F., Navarre, D.A. and Williamson, V.M. 2004. Salicylic acid is part of the Mi-1-mediated defense response to root-knot nematode in tomato. Mol Plant Microbe Interact. 17(4), pp.351–356.

Cecchini, N.M., Steffes, K., Schläppi, M.R., Gifford, A.N. and Greenberg, J.T., 2015. Arabidopsis AZI1 family proteins mediate signal mobilization for systemic defence priming. Nat Commun. 6(1), p.7658.

Conrath, U., Beckers, G.J., Flors, V., García-Agustín, P., Jakab, G., Mauch, F., Newman, M.A., Pieterse, C.M., Poinssot, B., Pozo, M.J. and Pugin, A. 2006. Priming: getting ready for battle. Mol Plant Microbe Interact. 19(10), pp.1062–1071.

Coyne, D., Smith, M. and Plowright, R. 2001. Plant parasitic nematode populations on upland and hydromorphic rice in Côte d’Ivoire: relationship with moisture availability and crop development on a valley slope. Agriculture, ecosystems & environment. 84(1), pp.31–43.

de Carvalho, L.M., Benda, N.D., Vaughan, M.M., Cabrera, A.R., Hung, K., Cox, T., Abdo, Z., Allen, L.H. and Teal, P.E. 2015. Mi-1-mediated nematode resistance in tomatoes is broken by short-term heat stress but recovers over time. J Nematol. 47(2), p.133.

Dhar, N., Caruana, J., Erdem, I. and Raina, R. 2020. An Arabidopsis DISEASE RELATED NONSPECIFIC LIPID TRANSFER PROTEIN 1 is required for resistance against various phytopathogens and tolerance to salt stress. Gene. 753, p.144802.

Dropkin, V.H. 1969. The necrotic reaction of tomatoes and other hosts resistant to Meloidogyne: reversal by temperature. Phytopathology. 59, pp.1632–1637.

Elmore, J.M. and Coaker, G. 2011. The role of the plasma membrane H+-ATPase in plant– microbe interactions. Mol Plant. 4(3), pp.416–427.

Escobar, C., Barcala, M., Cabrera, J. and Fenoll, C. 2015. Overview of root-knot nematodes and giant cells. In Advances in botanical research (Vol. 73, pp. 1-32). Academic Press.

Fasan, T. and Haverkort, A.J. 1991. The influence of cyst nematodes and drought on potato growth. 1. Effects on plant growth under semi-controlled conditions. Netherlands Journal of Plant Pathology. 97, pp.151–161.

García-Olmedo, F., Molina, A., Segura, A. and Moreno, M. 1995. The defensive role of nonspecific lipid-transfer proteins in plants. Trends Microbiol. 3(2), pp.72–74.

Grant, M. and Lamb, C. 2006. Systemic immunity. Curr Opin Plant Biol. 9(4), pp.414–420.

Haverkort, A.J. and Verhagen, A. 2008. Climate change and its repercussions for the potato supply chain. Potato research. 51, pp.223–237.

He, Y., Liu, Y., Li, M., Lamin-Samu, A.T., Yang, D., Yu, X., Izhar, M., Jan, I., Ali, M. and Lu, G. 2021. The Arabidopsis SMALL AUXIN UP RNA32 protein regulates ABA-mediated responses to drought stress. Front Plant Sci. 12, p.625493.

Ilyas, M., Nisar, M., Khan, N., Hazrat, A., Khan, A.H., Hayat, K., Fahad, S., Khan, A. and Ullah, A. 2021. Drought tolerance strategies in plants: a mechanistic approach. J Plant Growth Regul. 40, pp.926–944.

Jammes, F., Lecomte, P., de Almeida-Engler, J., Bitton, F., Martin-Magniette, M.L., Renou, J.P., Abad, P. and Favery, B. 2005. Genome-wide expression profiling of the host response to root-knot nematode infection in Arabidopsis a. Plant J. 44(3), pp.447–458.

Janiak, A., Kwaśniewski, M. and Szarejko, I. 2016. Gene expression regulation in roots under drought. J Exp Bot. 67(4), pp.1003–1014.

Jung, H.W., Tschaplinski, T.J., Wang, L., Glazebrook, J. and Greenberg, J.T. 2009. Priming in systemic plant immunity. Science. 324(5923), pp.89–91.

Kim, J.H., Castroverde, C.D.M., Huang, S., Li, C., Hilleary, R., Seroka, A., Sohrabi, R., Medina-Yerena, D., Huot, B., Wang, J. and Nomura, K. 2022. Increasing the resilience of plant immunity to a warming climate. Nature. 607(7918), pp.339–344.

Kirubakaran, S.I., Begum, S.M., Ulaganathan, K. and Sakthivel, N. 2008. Characterization of a new antifungal lipid transfer protein from wheat. Plant Physiol Biochem. 46(10), pp.918–927.

Kreye, C., Bouman, B.A.M., Reversat, G., Fernandez, L., Cruz, C.V., Elazegui, F., Faronilo, J.E. and Llorca, L. 2009. Biotic and abiotic causes of yield failure in tropical aerobic rice. Field Crops Research. 112(1), pp.97–106.

Lim, G.H., Shine, M.B., de Lorenzo, L., Yu, K., Cui, W., Navarre, D., Hunt, A.G., Lee, J.Y., Kachroo, A. and Kachroo, P. 2016. Plasmodesmata localizing proteins regulate transport and signaling during systemic acquired immunity in plants. Cell Host Microbe. 19(4), pp.541–549.

Liu, F., Zhang, X., Lu, C., Zeng, X., Li, Y., Fu, D. and Wu, G. 2015. Non-specific lipid transfer proteins in plants: presenting new advances and an integrated functional analysis. J Exp Bot. 66(19), pp.5663–5681.

Liu, J., Elmore, J.M., Fuglsang, A.T., Palmgren, M.G., Staskawicz, B.J. and Coaker, G. 2009. RIN4 functions with plasma membrane H+-ATPases to regulate stomatal apertures during pathogen attack. PLoS Biol. 7(6), p.e1000139.

Loake, G. and Grant, M. 2007. Salicylic acid in plant defence—the players and protagonists. Curr Opin Plant Biol. 10(5), pp.466–472.

Martin-StPaul, N., Delzon, S. and Cochard, H. 2017. Plant resistance to drought depends on timely stomatal closure. Ecology letters. 20(11), pp.1437–1447.

Mauch-Mani, B., Baccelli, I., Luna, E. and Flors, V. 2017. Defense priming: an adaptive part of induced resistance. Annu Rev Plant Biol. 68, pp.485–512.

Missaoui, K., Gonzalez-Klein, Z., Pazos-Castro, D., Hernandez-Ramirez, G., Garrido-Arandia, M., Brini, F., Diaz-Perales, A. and Tome-Amat, J. 2022. Plant non-specific lipid transfer proteins: An overview. Plant Physiol Biochem. 171, pp.115–127.

Mittler, R. 2006. Abiotic stress, the field environment and stress combination. Trends Plant Sci. 11(1), pp.15–19.

Molina, A. and García-Olmedo, F. 1993. Developmental and pathogen-induced expression of three barley genes encoding lipid transfer proteins. Plant J. 4(6), pp.983–991.

Muhammad Aslam, M., Waseem, M., Jakada, B.H., Okal, E.J., Lei, Z., Saqib, H.S.A., Yuan, W., Xu, W. and Zhang, Q. 2022. Mechanisms of abscisic acid-mediated drought stress responses in plants. Int J Mol Sci. 23(3), p.1084.

Nagpal, P., Reeves, P.H., Wong, J.H., Armengot, L., Chae, K., Rieveschl, N.B., Trinidad, B., Davidsdottir, V., Jain, P., Gray, W.M. and Jaillais, Y. 2022. SAUR63 stimulates cell growth at the plasma membrane. PLoS genetics. 18(9), p.e1010375.

Obermeyer, S., Stöckl, R., Schnekenburger, T., Moehle, C., Schwartz, U. and Grasser, K.D. 2022. Distinct role of subunits of the Arabidopsis RNA polymerase II elongation factor PAF1C in transcriptional reprogramming. Front Plant Sci. 13, p.974625.

Omae, N. and Tsuda, K. 2022. Plant-microbiota interactions in abiotic stress environments. Mol Plant Microbe Interact. 35(7), pp.511–526.

Pandey, P., Irulappan, V., Bagavathiannan, M.V. and Senthil-Kumar, M. 2017. Impact of combined abiotic and biotic stresses on plant growth and avenues for crop improvement by exploiting physio-morphological traits. Front Plant Sci. 8, p.537.

Parker, J.E. 2009. The quest for long-distance signals in plant systemic immunity. Sci Signal. 2(70), pp.pe31–pe31.

Prasch, C.M. and Sonnewald, U. 2013. Simultaneous application of heat, drought, and virus to Arabidopsis plants reveals significant shifts in signaling networks. Plant Physiol. 162(4), pp.1849–1866.

Qiu, T., Qi, M., Ding, X., Zheng, Y., Zhou, T., Chen, Y., Han, N., Zhu, M., Bian, H. and Wang, J. 2020. The SAUR41 subfamily of SMALL AUXIN UP RNA genes is abscisic acid inducible to modulate cell expansion and salt tolerance in Arabidopsis thaliana seedlings. Ann Bot. 125(5), pp.805–819.

Rasheed, S., Bashir, K., Matsui, A., Tanaka, M. and Seki, M. 2016. Transcriptomic analysis of soil-grown Arabidopsis thaliana roots and shoots in response to a drought stress. Front Plant Sci. 7, p.180.

Rasmussen, S., Barah, P., Suarez-Rodriguez, M.C., Bressendorff, S., Friis, P., Costantino, P., Bones, A.M., Nielsen, H.B. and Mundy, J. 2013. Transcriptome responses to combinations of stresses in Arabidopsis. Plant physiology. 161(4), pp.1783–1794.

Rivero, R.M., Mestre, T.C., Mittler, R., Rubio, F., Garcia-Sanchez, F. and Martinez, V. 2014. The combined effect of salinity and heat reveals a specific physiological, biochemical and molecular response in tomato plants. Plant Cell Environ. 37(5), pp.1059–1073.

Rizhsky, L., Liang, H. and Mittler, R. 2002. The combined effect of drought stress and heat shock on gene expression in tobacco. Plant Physiol. 130(3), pp.1143–1151.

Sinha, R., Zandalinas, S.I., Fichman, Y., Sen, S., Zeng, S., Gómez-Cadenas, A., Joshi, T., Fritschi, F.B. and Mittler, R. 2022. Differential regulation of flower transpiration during abiotic stress in annual plants. New Phytologist. 235(2), pp.611–629.

Sondergaard, T.E., Schulz, A. and Palmgren, M.G. 2004. Energization of transport processes in plants. Roles of the plasma membrane H+-ATPase. Plant Physiol. 136(1), pp.2475–2482.

Spartz, A.K., Ren, H., Park, M.Y., Grandt, K.N., Lee, S.H., Murphy, A.S., Sussman, M.R., Overvoorde, P.J. and Gray, W.M. 2014. SAUR inhibition of PP2C-D phosphatases activates plasma membrane H+-ATPases to promote cell expansion in Arabidopsis. Plant Cell. 26(5), pp.2129–2142.

Stortenbeker, N. and Bemer, M. 2019. The SAUR gene family: the plant’s toolbox for adaptation of growth and development. J Exp Bot. 70(1), pp.17–27.

Suzuki, N., Rivero, R.M., Shulaev, V., Blumwald, E. and Mittler, R. 2014. Abiotic and biotic stress combinations. New Phytol. 203(1), pp.32–43.

Taylor, S.C., Nadeau, K., Abbasi, M., Lachance, C., Nguyen, M. and Fenrich, J. (2019) The ultimate qPCR experiment: producing publication quality, reproducible data the first time. Trends Biotech. 37, 761–774.

Triky-Dotan, S., Yermiyahu, U., Katan, J. and Gamliel, A. 2005. Development of crown and root rot disease of tomato under irrigation with saline water. Phytopathology. 95(12), pp.1438–1444.

van Butselaar, T. and Van den Ackerveken, G. 2020. Salicylic acid steers the growth–immunity tradeoff. Trends Plant Sci. 25(6), pp.566–576.

Vos, I.A., Pieterse, C.M. and Van Wees, S.C. 2013. Costs and benefits of hormone-regulated plant defences. Plant Pathology. 62, pp.43–55.

Wildermuth, M.C., Dewdney, J., Wu, G. and Ausubel, F.M. 2001. Isochorismate synthase is required to synthesize salicylic acid for plant defence. Nature. 414(6863), pp.562–565.

Wong, J.H., Klejchová, M., Snipes, S.A., Nagpal, P., Bak, G., Wang, B., Dunlap, S., Park, M.Y., Kunkel, E.N., Trinidad, B. and Reed, J.W. 2021. SAUR proteins and PP2C. D phosphatases regulate H+-ATPases and K+ channels to control stomatal movements. Plant Physiol. 185(1), pp.256–273.

Yamaguchi, Y.L., Suzuki, R., Cabrera, J., Nakagami, S., Sagara, T., Ejima, C., Sano, R., Aoki, Y., Olmo, R., Kurata, T. and Obayashi, T. 2017. Root-knot and cyst nematodes activate procambium-associated genes in Arabidopsis roots. Front Plant Sci. 8, p.1195.

Zacheo, G. and Bleve-Zacheo, T. 1984. Influence of different temperatures on resistance of tomato plants to Meloidogyne incognita. Nematologia Mediterranea.

